# Phenotype integration improves power and preserves specificity in biobank-based genetic studies of MDD

**DOI:** 10.1101/2022.08.15.503980

**Authors:** Andrew Dahl, Michael Thompson, Ulzee An, Morten Krebs, Vivek Appadurai, Richard Border, Silviu-Alin Bacanu, Thomas Werge, Jonathan Flint, Andrew J. Schork, Sriram Sankararaman, Kenneth Kendler, Na Cai

## Abstract

Biobanks often contain several phenotypes relevant to a given disorder, and researchers face complex tradeoffs between shallow phenotypes (high sample size, low specificity and sensitivity) and deep phenotypes (low sample size, high specificity and sensitivity). Here, we study an extreme case: Major Depressive Disorder (MDD) in UK Biobank. Previous studies found that shallow and deep MDD phenotypes have qualitatively distinct genetic architectures, but it remains unclear which are optimal for scientific study or clinical prediction. We propose a new framework to get the best of both worlds by integrating together information across hundreds of MDD-relevant phenotypes. First, we use phenotype imputation to increase sample size for the deepest available MDD phenotype, which dramatically improves GWAS power (increases #loci ~10 fold) and PRS accuracy (increases R2 ~2 fold). Further, we show the genetic architecture of the imputed phenotype remains specific to MDD using genetic correlation, PRS prediction in external clinical cohorts, and a novel PRS-based pleiotropy metric. We also develop a complementary approach to improve specificity of GWAS on shallow MDD phenotypes by adjusting for phenome-wide PCs. Finally, we study phenotype integration at the level of GWAS summary statistics, which can increase GWAS and PRS power but introduces non-MDD-specific signals. Our work provides a simple and scalable recipe to improve genetic studies in large biobanks by combining the sample size of shallow phenotypes with the sensitivity and specificity of deep phenotypes.

## Introduction

Although Major Depressive Disorder (MDD) is the most common psychiatric disorder and the leading cause of disability worldwide, its causes are largely unknown. Despite the moderate familial heritability of MDD (~40%)^1^, genome-wide association studies (GWAS) have only recently begun to identify replicable risk loci and polygenic risk scores (PRS)^2–7^. These discoveries were enabled by increasing power along two primary dimensions: depth of phenotyping and sample size^8^. Increasing sample size improves GWAS and PRS power by reducing standard errors of estimated genetic effects on a given MDD phenotype^2,9^. Alternatively, increasing diagnostic accuracy through structured clinical interviews prevents dilution of genetic effect sizes, thus improving GWAS power^7,8,10^ and PRS accuracy^10,11^. In practice, studies have a fixed budget and must always tradeoff between increasing sample size or phenotyping depth. The optimal choice for current and future MDD studies remains contested^10,12,13^. Ultimately, it depends on our goals.

One important goal is statistical explanation, defined as the number of GWAS hits or the PRS prediction accuracy. Most MDD GWAS have focused on this goal, which is best achieved by maximizing sample size^10,11^. This motivates the use of shallow phenotypes in large biobanks, including self-reported depression or depression treatment^3,5^. Sample sizes are often further increased by using health records of seeking care for depression (e.g. iPSYCH^14^, Million Veterans Project^6^). This more accurately represents the population because it is not based on volunteering, though the accuracy and consistency of diagnostic criteria will vary among studies and study sites. These studies have amassed sample sizes of millions of individuals and have identified hundreds of risk loci, as well as PRS with state-of-the-art prediction accuracy in European-ancestry clinical cohorts^2–6^.

A partly distinct goal is biological insight. This is more difficult to measure or even define, but it represents one of the ultimate goals of genetics: characterizing biological mechanisms to improve prediction and treatment for all. This goal may never be achieved by increasing sample size with shallow phenotyping, because shallow phenotypes are confounded by genetic effects that do not pertain to MDD biology^10^. In contrast, deep phenotyping in clinical cohorts (e.g., PGC^2^, CONVERGE^7^) has identified a handful of replicated genetic loci that could potentially generate hypotheses on MDD-specific biology. However, this has not yet been demonstrated robustly, as current sample sizes simply do not provide power to yield enough genetic signals for definitive biological inferences^7^.

In this paper, we propose to bridge the shallow-deep gap by integrating information across hundreds of MDD-relevant phenotypes in UK Biobank^10,15^ (UKB, **Figure 1**). We focus on using phenotype imputation^16,17^ to increase the effective sample size for the deepest MDD phenotype in UKB (LifetimeMDD)^10^, which dramatically improves GWAS power and PRS accuracy over any individual MDD phenotype^18^. We extensively characterize the genetic architecture underlying these imputed phenotypes and show they remain specific to LifetimeMDD. Further, we develop a novel approach to partly remove non-specific signals from GWAS on shallow phenotypes akin to latent factor corrections in eQTL studies^19–22^. We also investigate phenotype integration via GWAS summary statistics using MTAG, which offers varying specificity and sensitivity depending on input choices. Finally, we developed a novel metric to quantify the specificity of a given PRS, which demonstrates that imputed deep phenotypes of MDD are both more specific and more sensitive than observed shallow phenotypes.

**Figure 1.**
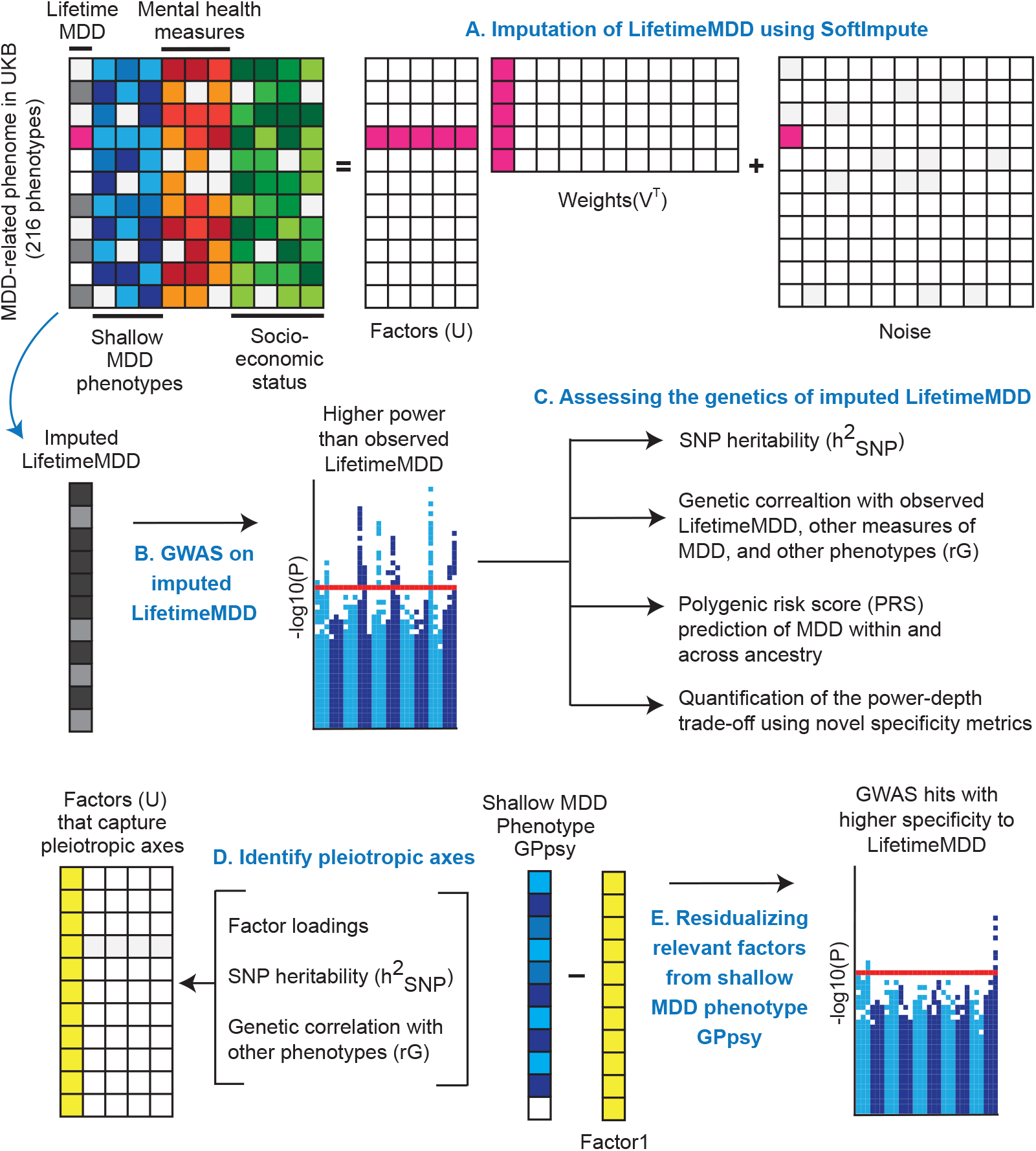
Study overview. **(A)** We impute LifetimeMDD using a partially-observed matrix of depression-relevant phenotypes in UK Biobank. We focus on using SoftImpute, which also produces latent phenome-wide factors. **(B)** We then perform GWAS on observed and imputed values of LifetimeMDD, as well as **(C)** downstream polygenic analyses, including in-sample and out-of-sample PRS predictions of MDD. We also study the genetic basis of the latent factors of the depression phenome **(D)**, and residualize latent factors from shallow MDD phenotypes to remove non-specific pleiotropic effects **(E)**.

## Results

### Phenotype imputation more than doubles effective sample size for LifetimeMDD

We focus on the deepest available measure of MDD in UKB^10^, LifetimeMDD, which we derive by applying clinical diagnostic criteria *in silico* to MDD symptom data from the PHQ-9 questionnaire and CIDI short form (CIDI-SF) in the online Mental Health Questionnaire. This procedure identifies 16,297 LifetimeMDD cases and 50,869 controls. Because most individuals did not complete these questionnaires, LifetimeMDD is missing for 269,962 individuals. We also study a shallow measure of MDD, GPpsy^10^, defined by seeking help from a General Practitioner for “depression, anxiety, tension, or nerves”. For imputation and downstream analyses, we use a broad depression-relevant phenome with 217 phenotypes, including comorbidities, family history, and socioeconomic, demographic, and environmental phenotypes (**Supplementary Note, Supplementary Table 1**).

We first impute the depression phenome using SoftImpute^23^ (**Methods**). We previously found SoftImpute to be the most scalable among several established approaches^16,23^. SoftImpute is a variant of principal component analysis (PCA) that accommodates missing data. It uses the observed phenotype data to identify latent factors, and then uses these factors to impute the missing data. As in our prior work, we tune SoftImpute’s regularization parameter using realistically held-out test data by taking unions of missingness patterns across samples^16^, and also use this approach to estimate the imputation accuracy for each phenotype (**Extended Data Figure 1**)^16^. Imputation accuracy varied widely across phenotypes, ranging from *R^2^*=1% for being a twin (1% missing) to *R^2^*=97% for neuroticism score (19% missing). For LifetimeMDD, we estimated the phenotype imputation *R^2^* to be 40% (80% missing). Roughly speaking, this means that SoftImpute more than doubles the effective sample size^16,24^ of LifetimeMDD (N observed=67K, N effective=166K, **Methods**). We show the distribution of imputed LifetimeMDD values in held-out cases and controls in **Extended Data Figure 1**. We found that the imputed measures had deflated variances and inflated correlations (**Supplementary Note**, **Supplementary Figure 1**), as expected^16^. This could bias some downstream tests, such as genetic correlations. One main goal in this work is to determine if this approach to phenotype imputation succeeds for large scale single-trait genetic studies.

As sex/gender significantly impacts MDD risk^25–27^, we repeated our SoftImpute analyses in each sex separately and compared the results to our primary approach (which includes sex as a column in the phenotype matrix). Overall, we find that the imputation *R^2^* are highly correlated across the female-, male-, and joint-imputation approaches (Pearson r between *R^2^* female and ß^2^joint = 0.928; between ß^2^male and ß^2^joint = 0.927; between ß^2^male and ß^2^female = 0.85, **Supplementary Figure 2**), without any statistically significant differences on average (average ß^2^female=21.7%, ß^2^male=20.1%, *R^2^* joint=21.3%, **Supplementary Table 2**).

Finally, we applied a new deep-learning imputation method, AutoComplete^28^, to the same phenotype matrix (**Methods**). AutoComplete improved estimated imputation accuracy for most phenotypes with >10% missingness (29/42), and increased average estimated *R^2^* by 2.9%.

### Phenotype imputation improves GWAS power for LifetimeMDD

We next assessed the impact of phenotype imputation on GWAS. We performed GWAS on observed LifetimeMDD (N=67,164), imputed values of LifetimeMDD (ImpOnly, N=269,962), and the concatenation of imputed and observed LifetimeMDD (ImpAll, N=337,126, **Methods**). GWAS on the observed values of LifetimeMDD identified one significant locus (**Figure 2E**). GWAS on the imputed values increased the number of GWAS loci to 13 and 18 for SoftImpute and AutoComplete, respectively (**Figure 2A,B, Supplementary Table 3**). Finally, GWAS on the combination of both imputed and observed values further increased the number of significant loci to 26 and 40 for SoftImpute and AutoComplete, respectively (**Figure 2C,D, Supplementary Table 3**). We confirmed that these improvements in the number of GWAS hits over the single hit from observed LifetimeMDD are very unlikely to result purely from chance (**Supplementary Figure 3**).

**Figure 2.**
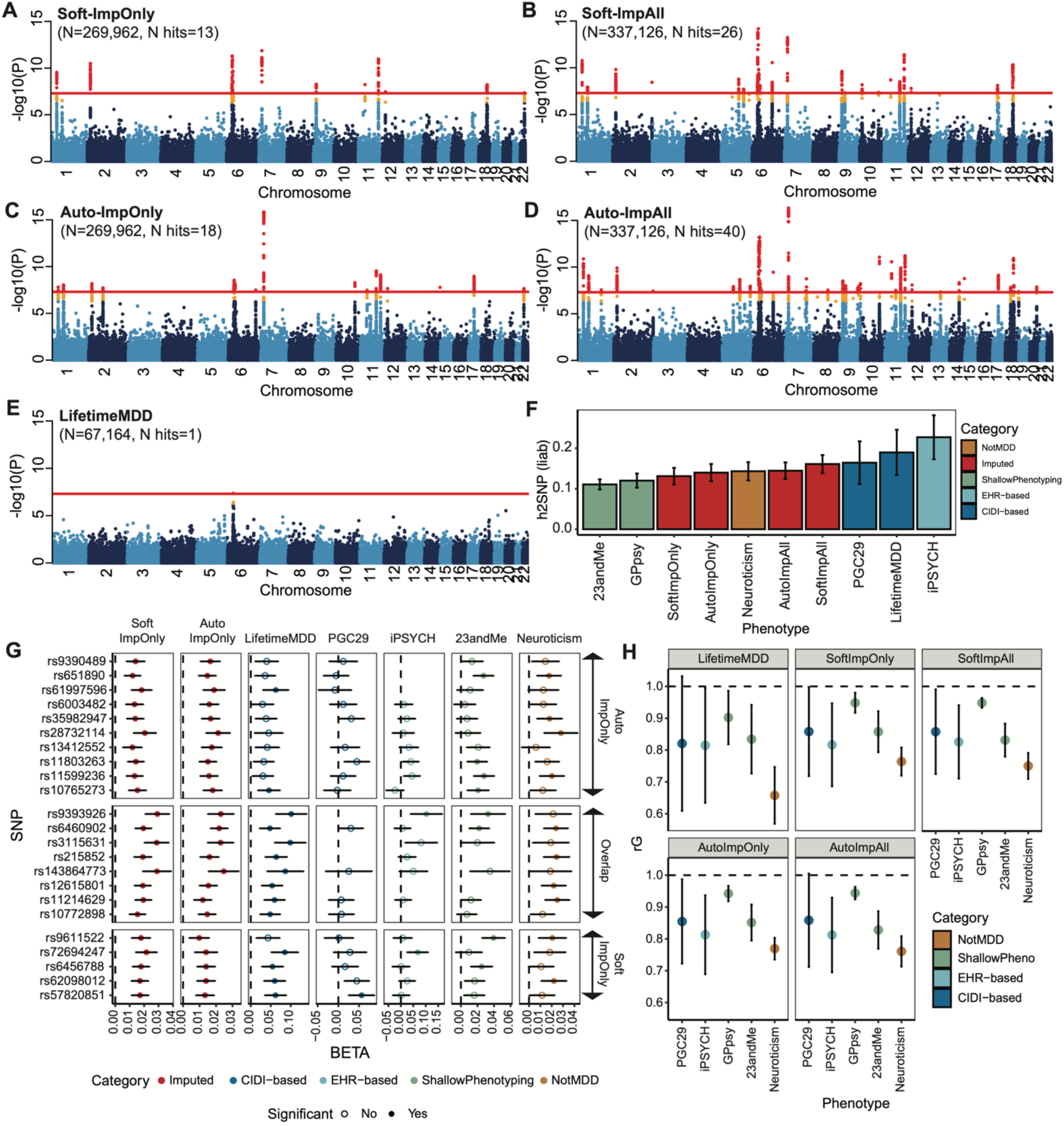
Genetic architecture of observed and imputed LifetimeMDD. Manhattan plots for GWAS on **(A,C)** imputed LifetimeMDD values from SoftImpute and AutoComplete (Soft-ImpOnly, Auto-ImpOnly, N=270K); **(B,D)** combined imputed and observed LifetimeMDD values from SoftImpute and AutoComplete (Soft-ImpAll, Auto-ImpAll, N=337K); and **(E)** observed LifetimeMDD (N=67K). Red lines show the genome-wide significance threshold of P < 5×10-8; **(F)** Liability-scale estimates of SNP-based heritability and **(H)** genetic correlation between all UKB measures of MDD and external MDD studies from PGC, iPSYCH, and 23andMe. **(G)** Replication of GWAS effect sizes from Soft-ImpOnly and Auto-ImpOnly in observed LifetimeMDD and external MDD studies. All error bars indicate 95% confidence intervals.

We investigated if the new GWAS hits from phenotype imputation were specific to MDD biology by comparing the ImpOnly GWAS to other MDD GWAS. First, we compared the two imputation methods. Out of 13 and 18 GWAS loci for ImpOnly from SoftImpute and AutoComplete respectively, 8 overlap (giving a total of 23, **Extended Data Figure 2**). Further, 9 of the remaining 15 loci had P < 10^-5^ in both ImpOnly GWAS and all 15 have P < 0.05/23. This shows our two imputation methods capture highly overlapping genetic signals, but AutoComplete has greater power. Next, we assessed these 8 shared hits in four non-overlapping depression cohorts (**Methods**, **Supplementary Note**): observed LifetimeMDD in UKB; self-reported depression diagnosis or treatment in 23andMe^5^; the 29 MDD cohorts of the Psychiatric Genomics Consortium^2^ (PGC29); and Danish registry data on MDD cases and population controls (iPSYCH^14,29^). For reference, we also compared to UKB measures of neuroticism, a personality trait that is genetically correlated but distinct from MDD^30^. We found that all 8 hits shared between both ImpOnly GWAS have sign-consistent effect size estimates across all of these depression cohorts, as well as neuroticism. Moreover, all 8 are significant for observed LifetimeMDD in UKB at P < 0.05/23. Finally, out of the 23 SNPs significant in one ImpOnly GWAS, 18 replicate in at least one GWAS of observed MDD at P < 0.05/23 (**Extended Data Figure 2**). Altogether, these results show that the predominant loci underlying imputed LifetimeMDD are relevant to the biology of MDD.

We then checked if the ImpOnly GWAS preserved the polygenic architecture of LifetimeMDD in terms of heritability and genetic correlation. First, we found that the liability scale SNP-based heritability (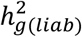 from LDSC^31^) was lower for imputed (Soft-ImpOnly 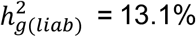 SE=1.0%; Auto-ImpOnly 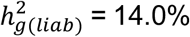, SE=1.1%) than observed LifetimeMDD (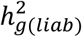, SE=2.9%, **Figure 2F**). This suggests that imputed values are noisier than observed LifetimeMDD. Nonetheless, the genetic correlations between imputed and observed LifetimeMDD are close to 1 (Soft-ImpOnly *r_g_* = .97, SE=.02; Auto-ImpOnly *r_g_* = .96, SE = .03), as are the correlations between imputed values from the two imputation methods (*r_g_* = 1.00, SE = .004). Moreover, the *r_g_* between ImpOnly phenotypes and secondary depression-related phenotypes largely mirror *r_g_* based on observed LifetimeMDD (**Figure 2G**). Altogether, imputed LifetimeMDD harbors similar genetic effects as observed LifetimeMDD, though it has additional sources of non-genetic noise.

Finally, we tested for effect size heterogeneity between the ImpOnly and observed LifetimeMDD GWAS. We used a simple random effect meta-analysis^32^ (**Methods**), as ImpOnly and observed LifetimeMDD GWAS use non-overlapping individuals. We find no significant heterogeneity between ImpOnly and observed LifetimeMDD at genome-wide significance (**Extended Data Figure 2**), and across the 13 and 18 GWAS hits in Soft-ImpOnly and Auto-ImpOnly, respectively, 6 and 4 SNPs showed significant heterogeneity at P < 0.05/23. Altogether, imputed LifetimeMDD has more non-genetic noise than observed LifetimeMDD, but has similar genetic architecture.

### Phenome-wide factors partition pleiotropic axes of depression risk

In order to understand the phenotypic correlations driving imputation, we examined the top latent factors in SoftImpute. These latent factors are essentially PCs of our depression-relevant phenome. We used two statistical metrics to prioritize factors for genetic study. First, we quantified the variance explained by each factor (**Methods**, **Figure 3A**). The top handful of factors clearly stood out from the background, but factors became comparable to background noise levels around factor 30. Second, we quantified factor stability by calculating the *R^2^* between factors estimated on separate halves of the data (**Methods, Figure 3B**). This is a variant of the prediction strength metric for clustering^33^. We found that the first 10 factors were extremely stable (min *R^2^*~98%), with stability decaying steadily afterward (factors 11-20 have average *R^2^*~80%, and 21-30 have average *R^2^*~60%). Overall, we conclude that the first ten or so factors are statistically meaningful.

**Figure 3.**
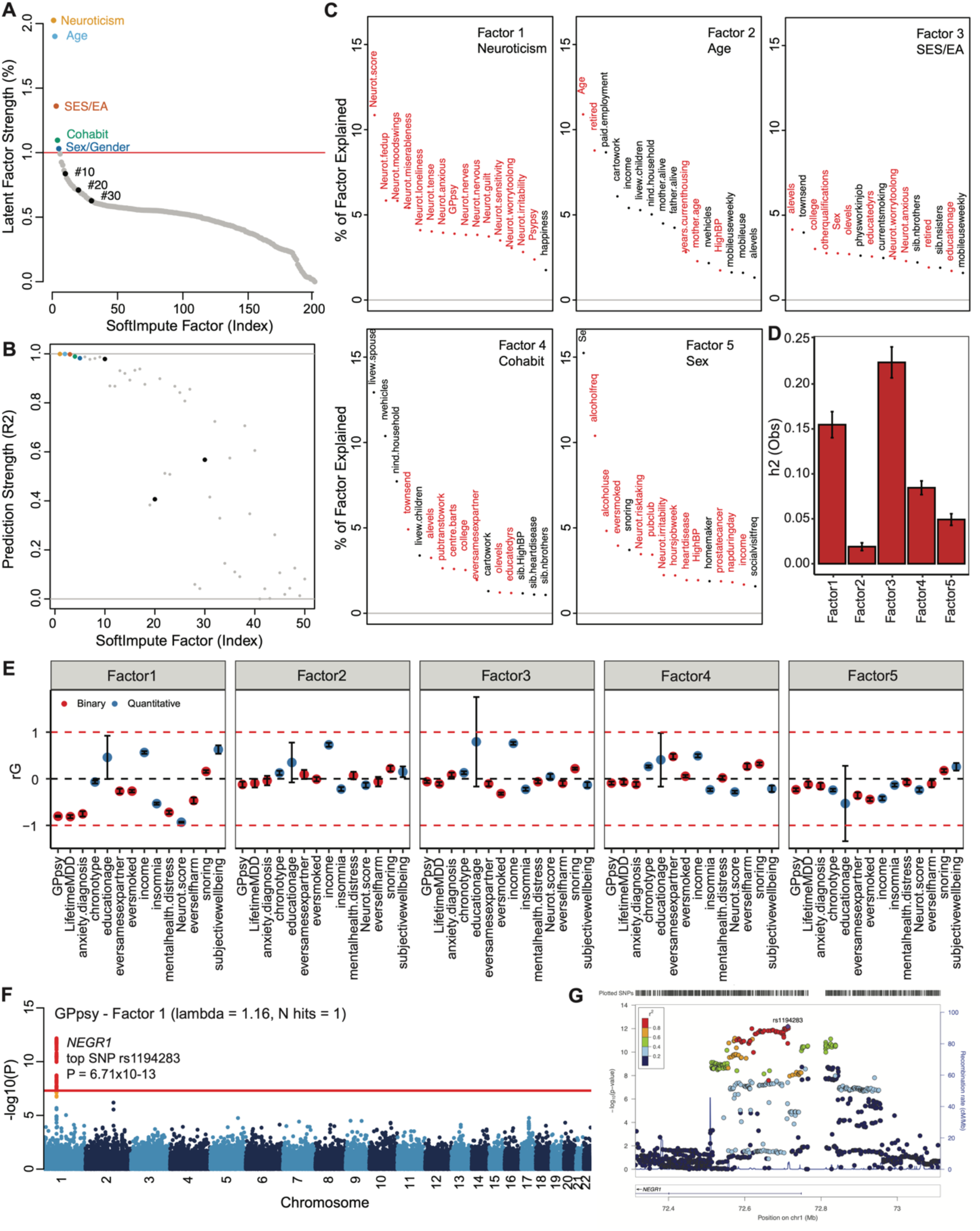
Characterizing top latent factors driving SoftImpute. Statistical importance of each factor measured by **(A)** percentage variance explained in the phenotype matrix and **(B)** factor prediction strength. **(C)** Top phenotype loadings for the top 5 Softmpute factors. **(D)** Estimates of heritability and **(E)** genetic correlations of the top 5 SoftImpute factors to MDD-relevant traits. **(F)** GWAS Manhattan plot of GPpsy conditioning on SoftImpute Factor 1; red line shows the genome-wide significance threshold. **(G)** Locus-zoom plot of the significant GWAS locus on gene *NEGR1*. All error bars indicate 95% confidence intervals.

Conservatively, we interpret only the top five factors. We name Factor 1 Neuroticism: its top loading is total neuroticism score, and it heavily loads on specific neuroticism items and shallow depression phenotypes^10^ (**Figure 3C, Supplementary Table 1)**. Factor 2 (Age) captures age and related socioeconomic variables, like retirement. Factor 3 (SES/EA) reflects complex socioeconomic status (SES) phenotypes, particularly education attainment (EA) and Townsend deprivation index. Factor 4 (Cohabit) is another complex social dimension, loading primarily on cohabitation phenotypes. Finally, Factor 5 (Sex/Gender) reflects sex/gender and known psychosocial correlates such as alcohol and tobacco use. Deeper factors are shown in **Supplementary Figure 4**.

We then studied the genetic basis of each factor with GWAS. Each factor had GWAS hits, ranging from from 3 (Age) to 309 (SES/EA), with *λ_GC_* ranging from 1.15 (Age) to 2.11 (SES/EA) (**Supplementary Figure 5**). We next estimated heritability for each factor and found that they range from 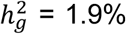 (SE = 0.2%) for Age to 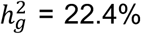 (SE=0.9%) for SES/EA (**Figure 3D, Supplementary Figure 4**). These results are consistent with our interpretations based on the factor loadings: Age has low 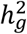 and few GWAS hits, while Neuroticism and SES/EA have high 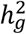, high *λ_GC_*, and many more GWAS hits. Finally, we profiled the genetic correlation between factors and various MDD phenotypes and related phenotypes (**Figure 3E, Supplementary Figure 4**). We found that the *r_g_* closely mirrored the factor loadings, which are based only on phenotypic correlations. For example, Factor 1 had *r_g_*= −0.93 (SE = 0.01) with neuroticism, and SES/EA had *r_g_*”=0.79 (SE = 0.96) with years of education and *r_g_*=0.75 (SE = 0.03) with income.

Given these results, we hypothesized that our top phenome-wide factors partly capture the nonspecific pathways that contribute to shallow MDD phenotypes. To test this hypothesis, we performed GWAS on a shallow MDD measure, GPpsy (N=332,629), conditioning on Factor 1. This is akin to removing confounders like batch effects in eQTL studies through conditioning on latent factors. We found that only 1 of the 25 GWAS hits for GPpsy remains after adjusting for Factor 1 (**Figure 3F**). This hit overlaps the gene *NEGR1* (top SNP rs1194283, OR = 1.05, SE = 0.0065, P = 6.71×10^-13^, **Figure 3G**), which has been identified as an MDD risk locus in multiple GWAS studies with varying phenotyping approaches^2,4,6,34–36^. Intriguingly, this locus also has replicated associations with body mass index and obesity in diverse populations^37–40^, suggesting it may act on MDD through a metabolic pathway that is independent of neuroticism. We also tested adjusting for each of the other top 10 factors. Generally, these adjustments had little impact, and only removed one or a few GWAS hits (**Supplementary Figure 6**). One clear exception, however, was adjustment for the SES/EA factor, which increased the number of GPpsy GWAS hits from 25 to 35. While this may seem surprising, false positives are expected after adjusting for a heritable latent factor^22,41,42^.

### MTAG is sensitive to inputs but improves GWAS power

As an alternative to phenotype imputation, we next evaluated phenotype integration at the summary statistic level via MTAG (multi-trait analysis of GWAS^18^), an inverse-covariance-weighted meta-analysis for GWAS on multiple traits. We did not use all 217 phenotypes in MTAG for two reasons. First, MTAG requires running GWAS on each input phenotype, which is computationally intractable for hundreds of phenotypes. Second, MTAG accrues false positives as the number of traits grows^18^. Instead, we performed MTAG on 6 different sets of input phenotypes, each of which produces an integrated LifetimeMDD GWAS (**Figure 4A, Extended Data Figure 3, Supplementary Table 4, Supplementary Note**).

**Figure 4.**
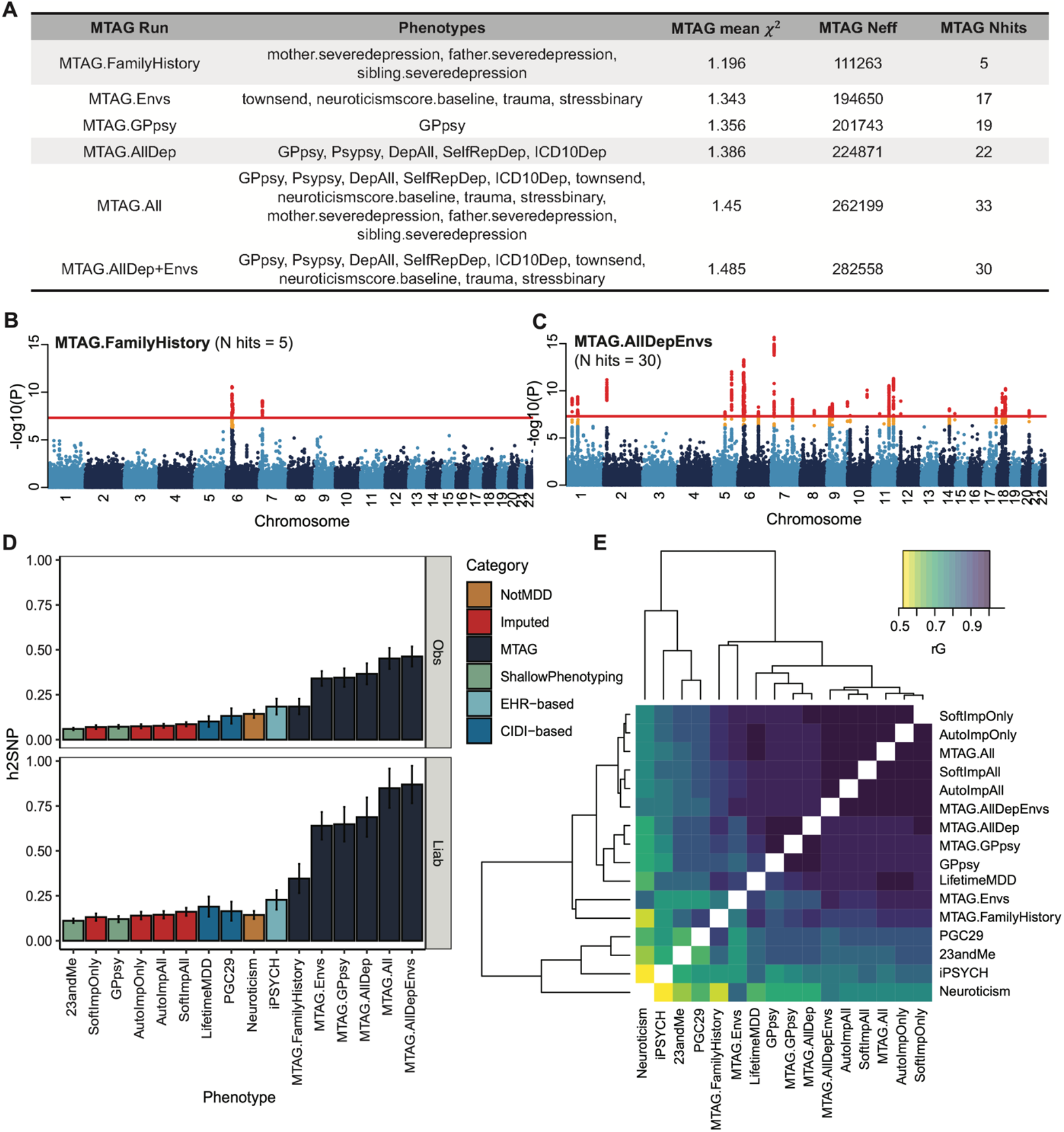
MTAG results for different choices of input phenotypes. **(A)** Description of the evaluated input choices for MTAG and their resulting GWAS summaries. The MTAG effective sample size refers to the power-equivalent sample sizes of MTAG GWAS vs single-trait GWAS^18^, calculated as N_eff_ = N_single_ * (*χ^2^*_MTAG_ - 1)/(*χ^2^*_single_ - 1), where *χ^2^* are the average GWAS chi-squared values. **(B,C)** Manhattan plots for MTAG models with fewest (MTAG.FamilyHistory) and greatest (MTAG.AllDep+Envs) number of GWAS hits; red line shows the genome-wide significance threshold. **(D)** SNP-based heritability estimates on the observed and liability scales for observed, imputed, and MTAG GWAS on LifetimeMDD as well as reference phenotypes. **(E)** Estimated genetic correlations for observed, imputed and MTAG analyses of LifetimeMDD and reference phenotypes. All error bars indicate 95% confidence intervals.

All MTAG input choices increased the number of GWAS hits from LifetimeMDD. On the low end, MTAG using family history measures of depression yielded 5 GWAS hits (MTAG.FamilyHistory, *λ_GC_*=1.20, **Figure 4B**). On the high end, MTAG using shallow MDD phenotypes and environmental factors (such as recent stressful life events, lifetime traumatic experiences, and townsend deprivation index) yielded 33 GWAS hits (MTAG.All, *λ_GC_*=1.45, **Figure 4C**). We note that MTAG.AllDep is analogous to depression phenotypes derived by manually combining similar input phenotypes^13^ and that MTAG.FamilyHistory is analogous to prior approaches that integrate family history measures into GWAS^43,44^. Of the total 51 hits across all MTAG runs, 34 overlap hits from the imputed GWAS with SoftImpute or AutoComplete (**Extended Data Figure 3**). Notably, we found that including more input phenotypes in MTAG always increased the number of GWAS hits. This is due to a combination of increased power to detect pleiotropic signals and increased false positive inflation^18^. Consistent with the latter contribution, MTAG GWAS yielded substantially inflated heritability estimates (on both the liability and observed scales), which increased with more input phenotypes. For example, MTAG.All gave 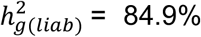 (SE = 5.6%), compared to 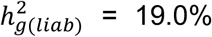 (SE = 2.9%) for observed LifetimeMDD (**Figure 4D, Supplementary Table 5**).

We next examined genetic correlations between MTAG and other MDD GWAS (**Figure 4E**). First, MTAG.All, which included the most input phenotypes, clustered together with the imputed GWAS, which leveraged all 217 phenotypes. Second, MTAG using shallow MDD phenotypes as input (MTAG.AllDep and MTAG.GPpsy) clustered with GWAS on GPpsy. Third, neuroticism is significantly more genetically correlated with MTAG.Envs (*i_g_* = 0.84, SE = 0.01) than LifetimeMDD (*r_g_* = 0.66, SE = 0.06). These results are consistent with prior observations that MTAG-based summary statistics modestly inflate genetic correlation to the input phenotypes^45^. Overall, the genetic correlations between MTAG and LifetimeMDD were high, with lowest value given by MTAG.Envs (*r_g_* = 0.90, SE = 0.03). Altogether, MTAG outputs resemble the chosen input phenotypes, and the choice of inputs significantly impacts power and specificity.

Finally, to compare like-to-like, we repeated our evaluation of SoftImpute imputation accuracy after restricting to the MTAG.All input phenotypes (and sex, age and 20 PCs). We found that imputation performed much worse with this reduced set of phenotypes (**Supplementary Figure 2**). For LifetimeMDD, specifically, imputation R2 dropped from 59.6% to 39.5% (P < 2×10-5, pooled t-test across folds). Overall, average imputation R2 on MTAG.All phenotypes dropped from 34.8% to 20.3% (**Supplementary Table 2**).

### Phenotype imputation and MTAG improves PRS accuracy for LifetimeMDD

We then assessed within-sample prediction accuracy of polygenic risk scores (PRS) based on integrated MDD phenotypes. We used 10-fold cross-validation to estimate the Nagelkerke’s *R^2^* prediction accuracy for LifetimeMDD in white British individuals in UKB. We jointly cross-validated the phenotype imputation and PRS construction (**Methods**). For MTAG, we jointly cross-validated the GWAS on secondary input phenotypes. To put these results in context, we compared to PRS built from observed LifetimeMDD (N = 67,164, Neff = 49,718) and GPpsy (N = 332,629, Neff = 298,992) in UKB^10^; MDD defined by structured interviews in PGC29^2^ (N = 42,455, Neff = 40,627); affective disorder defined by Danish health registries in iPSYCH^14^ (N = 38,123, Neff = 38,100) and self-reported depression in 23andMe^5^ (N = 307,354, Neff = 228,033, **Supplementary Table 6**).

We found imputing LifetimeMDD doubled PRS prediction accuracy over observed LifetimeMDD (**Figure 5A**, LifetimeMDD *R^2^* = 1.0%, 95% CI = [0.6%,1.4%], Soft-ImpAll *R^2^* = 2.1%, 95% CI = [1.3%, 2.9%], Auto-ImpAll *R^2^* = 2.2%, 95%CI=[1.4%,3.0%]). Consistent with prior reports^10,11^, we found that the GPpsy PRS predicts LifetimeMDD better than the LifetimeMDD PRS itself (*R^2^*=1.6%, 95% CI = [0.6%, 2.4%]), which is because GPpsy has roughly four times the sample size. Nonetheless, both SoftImpute and AutoComplete PRS outperformed the GPpsy PRS, demonstrating that integrating shallow and deep phenotypes can improve PRS over either alone. Finally, we found that the imputed LifetimeMDD PRS substantially outperform the PRS from iPSYCH (*R^2^* = 0.6%, 95% CI = [0.2%,0.9%]) and 23andMe (*R^2^* = 1.3%, 95% CI = [0.7%,1.9%]), even though iPSYCH used deeper phenotypes and 23andMe had a large sample size.

**Figure 5.**
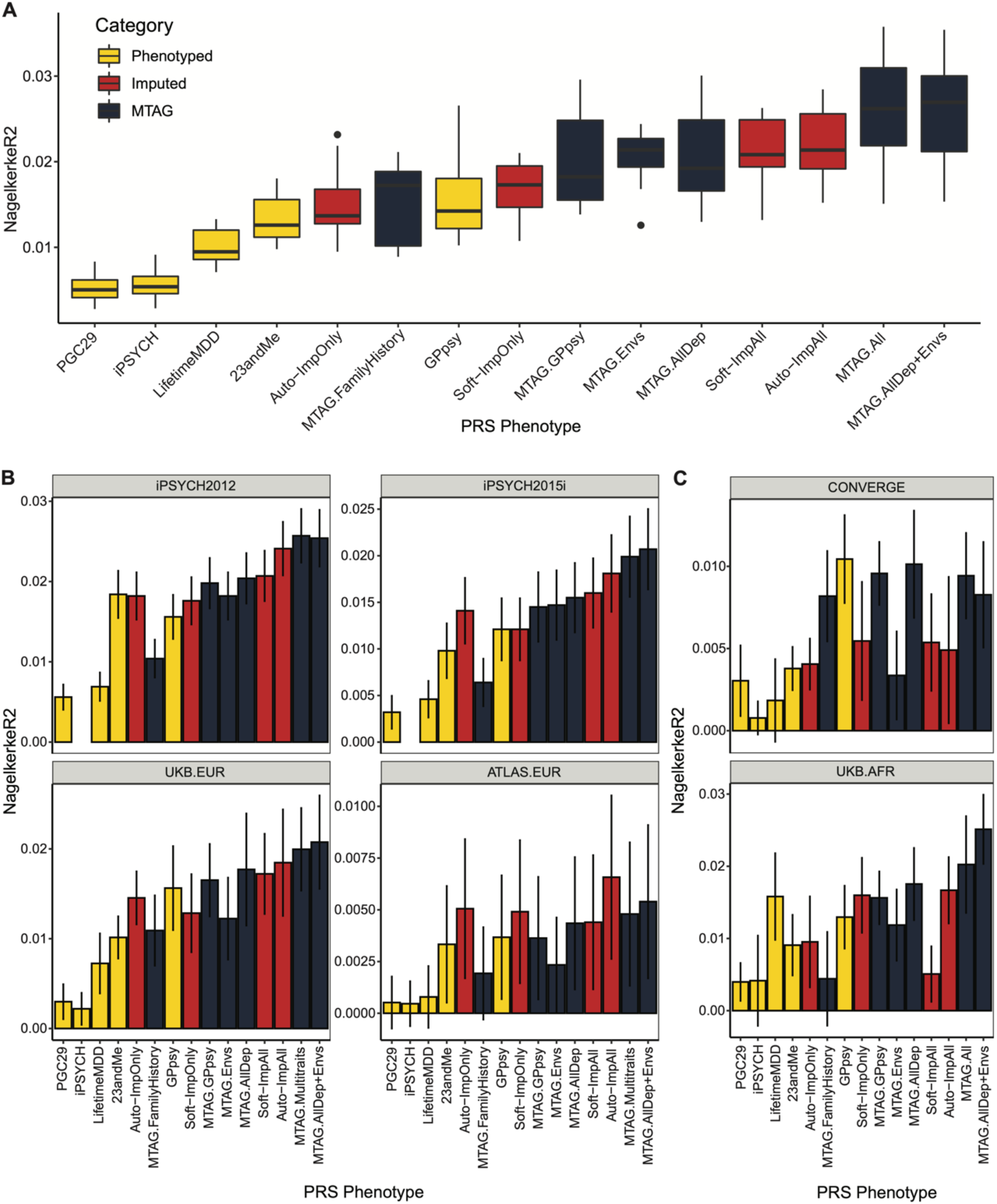
PRS performance using observed, imputed, and/or meta-analyzed MDD. **(A)** PRS prediction accuracy in the training population of unrelated white British individuals in UKB using 10-fold cross-validation. For imputed PRS, we also cross-validate the imputation. Median values are shown as a line in the box; whiskers of boxplots are 1.5 interquartile range; outliers outside of interquartile range are shown as filled dots. **(B)** Out-of-sample PRS prediction accuracy in four additional cohorts with European ancestries. **(C)** PRS prediction accuracy in African ancestry individuals in UKB and Han Chinese ancestry individuals in CONVERGE. All error bars indicate 95% confidence intervals.

The performance of MTAG PRS depended on the input phenotypes, mirroring the MTAG GWAS results (**Figure 4, Extended Data Figure 3**). For instance, MTAG.FamilyHistory does not substantially improve GWAS power, and its PRS underperforms imputed PRS (*R^2^* = 1.5%, 95% CI = [0.6%,2.5%], **Figure 5A**). On the other hand, MTAG.All significantly improves GWAS power and yields PRS that outperforms the imputed PRS by about 20% (*R^2^* = 2.6%, 95% CI =[1.3%,3.9%], **Figure 5A**). In particular, this demonstrates that MTAG with large numbers of inputs, which is non-standard and likely yields miscalibrated GWAS results, can nonetheless significantly improve PRS.

### Phenotype imputation and MTAG improves PRS portability

Having shown that phenotype integration improves PRS predictions in held-out white British individuals in UKB, we next asked if it also improves PRS predictions in different cohorts, diagnostic systems, and/or populations. Demonstrating portability is essential in order to establish that phenotype integration does not merely reflect dataset-specific biases. Measuring portability is also essential for assessing the clinical potential of PRS based on phenotype integration.

First, we tested PRS accuracy in non-British individuals with European ancestry in UKB (UKB.EUR, N=10,166). These individuals are measured on the same LifetimeMDD phenotype as our sample of white British UKB individuals and also have European ancestry, hence represent the most similar cohort (**Supplementary Note, Supplementary Figure 7**). Though the small sample size limits definitive conclusions, we observe a nearly identical pattern amongst PRS methods as in our training sample: imputation and MTAG almost always improve over both LifetimeMDD and GPpsy (**Figure 5B**). We next assessed portability to two large European-ancestry cohorts from iPSYCH (2012 cohort [N=42,250] and 2015i cohort [N=23,351], **Supplementary Note**). These non-overlapping samples are drawn from a nation-wide Danish birth cohort with diagnoses obtained from national health registers^29,46^. We again found qualitatively identical results, with imputation outperforming both LifetimeMDD and GPpsy, and the best MTAG setting outperforming imputation (**Figure 5B**). Finally, we tested portability to European-ancestry individuals in the ATLAS dataset based on MDD as defined in the UCLA EHR data^47,48^ (ATLAS.EUR, N=14,388, **Supplementary Note**, **Supplementary Figure 8, Supplementary Tables 6-8**). Again, small sample size prevents definitive comparisons, but phenotype imputation and the best MTAG setting improve estimated accuracy (**Figure 5B**).

We next tested these PRS in individuals with non-European genetic ancestries, including African ancestry individuals^5^ in UKB with observed LifetimeMDD status (UKB.AFR, N=687), as well as Han Chinese ancestry individuals in the CONVERGE cohort^7,49^ (N=10,502, **Supplementary Note**) who were assessed for severe, recurrent MDD (**Figure 5C**, **Supplementary Note, Supplementary Table 6**). Consistent with previous studies^50–52^, we find that the PRS we derived from GWAS on European-ancestry cohorts generally had poorer portability to non-European cohorts. Moreover, the strikingly consistent pattern of relative PRS accuracies observed in external European-ancestry cohorts no longer holds so clearly. In particular, it is surprising that the shallow PRS (using GPpsy) performs best in CONVERGE, which uses the deepest phenotyping of cohorts we study. Nonetheless, the best MTAG setting is always near-optimal, and PRS based on imputed LifetimeMDD always outperform PRS based on observed LifetimeMDD. Finally, we also tested PRS prediction accuracy in UKB individuals with Asian ancestry (UKB.ASN, N=334) as well as ATLAS individuals who self-identify as Latino (ATLAS.LAT, N=2,454), Black (ATLAS.AFR, N=1,158) or Asian (ATLAS.ASN, N=1,996). However, power was too low in these small cohorts for meaningful interpretation (**Supplementary Figure 9**).

### A new metric contrasts specificity of PRS from deep, shallow, and integrated phenotypes

While phenotype integration improves PRS prediction in UKB and in external cohorts, this may come at the cost of reduced specificity to MDD. This is because integration explicitly borrows information from secondary phenotypes, which could introduce genetic signals that are irrelevant to MDD. To quantify this spillover of non-specific effects into an MDD PRS, we compare prediction accuracy for LifetimeMDD to prediction accuracy for secondary phenotypes. We call this metric of specificity PRS Pleiotropy (*R^2^*_secondary_/*R^2^*_LifetimeMDD_).

Because core MDD biology is likely partly pleiotropic, its PRS Pleiotropy should be nonzero for many secondary phenotypes. We further expect that shallow MDD phenotypes, such as GPpsy, would generally have higher PRS Pleiotropy than LifetimeMDD^10^. Our main question, however, is whether phenotype integration suffers worse PRS Pleiotropy than shallow MDD phenotypes. For all PRS based on observed and integrated MDD, we calculated PRS Pleiotropy for 172 secondary phenotypes used in imputation. We then limited our investigation to the 62 secondary phenotypes that are significantly predicted by any examined PRS (P<0.05/172, **Methods**). Visualizing PRS Pleiotropy for observed LifetimeMDD across this depression phenome shows a spectrum of highly-linked traits, including shallow MDD phenotypes like GPpsy and genetically correlated traits like neuroticism, that quickly fades across successive phenotypes (**Figure 6A**). By comparison, GPpsy broadly has higher PRS Pleiotropy across secondary phenotypes, indicating GPpsy captures less-specific biology than LifetimeMDD, as expected. We also found that the 23andMe GWAS had similar PRS pleiotropy to GPpsy, consistent with the fact that both measure MDD by self-reported depression.

**Figure 6.**
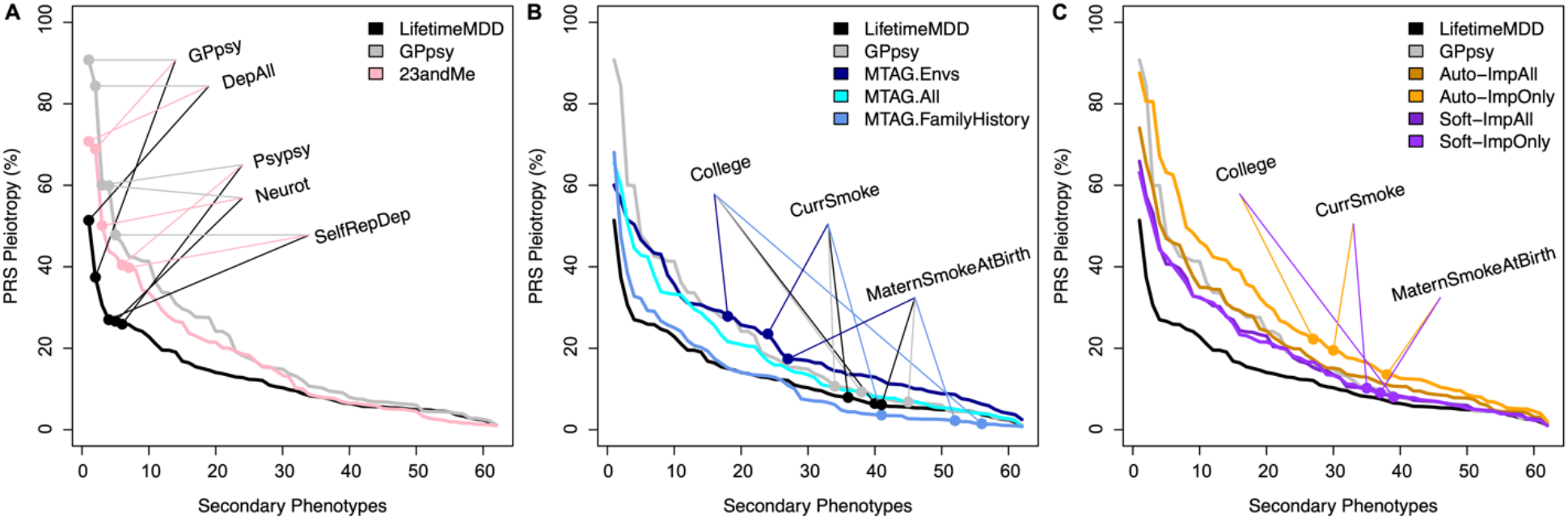
Phenome-wide PRS Pleiotropy quantifies non-specificity. PRS Pleiotropy spectra across the depression-relevant phenome, defined as the ratio of PRS prediction accuracy for secondary traits relative to LifetimeMDD (PRS Pleiotropy:= R^2^ _secondary_/R^2^ _LifetimeMDD_). **(A)** The PRS derived from GWAS on shallow MDD phenotypes (GPpsy or 23andMe) are less specific to LifetimeMDD than the PRS derived from GWAS on LifetimeMDD. **(B)** MTAG-based PRS range from highly specific (MTAG.FamilyHistory) to less specific than shallow MDD phenotypes (MTAG.Envs). **(C)** Softimpute PRS are more specific than the shallow PRS, while Autocomplete PRS are similar. Note that GPpsy and LifetimeMDD are each used in two ways: To build the PRS, and to evaluate PRS Pleiotropy.

We next evaluated PRS Pleiotropy for MTAG and found that specificity depends highly on input phenotypes (**Figure 6B**). First, MTAG.Envs has far higher PRS Pleiotropy across than GPpsy, showing its improved prediction power over GPpsy comes at a cost in specificity. On the other hand, MTAG.All is similar to GPpsy in specificity and almost doubles PRS prediction accuracy for LifetimeMDD, hence MTAG.All is clearly superior to GPpsy. Finally, MTAG.FamilyHistory has the opposite properties: while it only modestly improves PRS prediction accuracy over observed LifetimeMDD, this benefit comes without loss of specificity. We then evaluated PRS Pleiotropy for imputed phenotypes **(Figure 6C)**. The SoftImpute ImpAll and ImpOnly PRS are both more specific to LifetimeMDD than the GPpsy PRS, which is remarkable given that imputed values are constructed from more than 200 phenotypes, including GPpsy. The AutoComplete ImpOnly PRS was less specific than GPpsy, on the other hand, though the ImpAll PRS was comparable.

Finally, we asked which secondary phenotypes had non-specific effects in excess of pleiotropy expected from core MDD biology. We define Excess PRS Pleiotropy as PRS Pleiotropy minus the PRS Pleiotropy of observed LifetimeMDD, which represents our best proxy for core MDD biology^10^. As expected, self-reported depression (GPpsy and 23andMe) had Excess Pleiotropy for most secondary traits, especially shallow MDD measures (**Extended Data Figure 4A**). Likewise, MTAG.Envs had substantial Excess PRS Pleiotropy, especially for socioeconomic measures like education years (**Extended Data Figure 4B**). Notably, MTAG.FamilyHistory had far less Excess PRS Pleiotropy than other MTAG settings or GPpsy, and in particular actually has less pleiotropy for the socioeconomic measures driving Excess PRS Pleiotropy for MTAG.Envs. Finally, SoftImp-All had lower Excess PRS Pleiotropy than GPPsy (41/62 phenotypes); however, AutoImp-All had higher Excess PRS Pleiotropy (**Extended Data Figure 4C**). Overall, MTAG choices can outperform imputation in PRS sensitivity or specificity, but imputation provides a simple, scalable, and robust approach that simultaneously achieves near-optimal power and specificity.

Because of the complex relationship between effective sample size, p-value threshold, and PRS *R^2^*, we evaluated PRS Pleiotropy after downsampling the GWAS used to build the PRS (**Supplementary Note**). Overall, we found that PRS Pleiotropy is stable, though it can be upwardly biased for sample sizes below 100K (which is likely due to low power). In particular, our results are robust to the slight differences in the training PRS sample sizes, though observed LifetimeMDD PRS Pleiotropy is a conservative baseline because it is trained on 67K individuals (**Extended Data Figure 5, Supplementary Figures 10-11)**. Further, we confirmed that these results persist when we use exactly the same SNPs in each PRS (**Supplementary Note**, **Extended Data Figure 6**).

## Discussion

In this paper we address the power-specificity tradeoff between deep and shallow MDD phenotypes by integrating them together using phenotype imputation or MTAG. We show that the integrated MDD phenotypes greatly improve GWAS power and PRS accuracy while, crucially, preserving the genetic architecture of MDD. We propose a novel metric to assess the disorder-specificity of a PRS that is widely applicable to biobank-based GWAS. This metric characterizes a power-specificity tradeoff. For MTAG, adding more phenotypes significantly increases power but can sacrifice specificity. Imputing LifetimeMDD with SoftImpute, on the other hand, preserves more specificity. Overall, our results demonstrate that both approaches to phenotype integration outperform either deep or shallow phenotypes alone, and that phenotype integration is a practical way to improve biobank-based GWAS.

Phenotype imputation is a simple and scalable approach that should be considered for most biobank-based genetic studies. A particularly important but challenging future application for phenotype imputation will be to longitudinal data, which is often sporadically measured across time points and individuals, especially in biobank data. This requires carefully modeling nonrandom missingness, which is expected to be severe in longitudinal data. One limitation of our specific imputation approaches is that they distort higher-order moments (**Supplementary Figure 1**), which will bias some downstream analyses, like genetic correlation. As such, it is essential to thoroughly validate results with external data. In future work, this could be addressed with multiple imputation^53^ or with downstream tests that allow different effect sizes or noise variances between imputed and observed phenotypes^54^.

MTAG has several important strengths and complements imputation. First, it operates at the summary statistics level, which enables incorporating external GWAS results and is far more computationally efficient than phenotype imputation once GWAS have been performed. Second, we found that MTAG.All generally outperforms imputation in terms of GWAS hits and PRS power and portability. However, the tradeoff is that MTAG generally has less specificity. A striking exception is MTAG.FamHist, which preserves specificity. This extends previous observations that careful methods can exploit family history to improve genetic studies^43,44,55^. However, MTAG is highly sensitive to input phenotype selection, and therefore requires extensive domain knowledge, just like previous approaches including combining multiple depression measures^13^ and GWAS by proxy^43^. There are several natural extensions to further improve the performance of MTAG for phenotype integration. First, we could incorporate local estimates of genetic correlation in MTAG, which have been used to improve LifetimeMDD PRS in UKB^56^. Second, we could directly combine PRS for multiple traits using weights that optimize prediction^57–59^. Third, we could develop a more systematic approach to choose MTAG inputs, which is important because this choice heavily impacts power and specificity. However, this search is limited by the computational cost of performing cross-validated GWAS on each considered trait. Fourth, parametric models of confounding in summary statistics, like GWAS-by-subtraction^60^, could improve specificity as well as power. However, these models rely on choosing appropriate inputs and causal models, neither of which is straightforward for heterogeneous disorders. Finally, we could use GWAS on imputed phenotypes as inputs for MTAG; however, this may exacerbate biases in imputation because MTAG leverages correlations between traits, which are biased by most imputation approaches (**Supplementary Figure 1**).

Our study has implications for improving disorder specificity in future MDD studies. We have worked on the deepest MDD phenotype in UKB, LifetimeMDD, which is derived by applying DSM-5 criteria *in silico* to self-rated MDD symptoms in the MHQ. This is in fact shallow compared to a clinical diagnosis based on a structured in-person interview, especially due to self-report biases and misdiagnoses that have been repeatedly demonstrated^61–64^. In the future, this bias could be mitigated using methods based on probability weights, which have been recently developed for GWAS applications^65,66^. However, it is also previously shown that LifetimeMDD lies on essentially the same genetic liability continuum as gold-standard MDD^10^. In particular, genetic effects on LifetimeMDD are likely to represent MDD-specific biology, hence they provide a reliable benchmark when interrogating the specificity of genetic effects on integrated phenotypes. More broadly, we acknowledge that the DSM-5 criteria for MDD themselves have significant shortcomings in reliability^65–67^. Nonetheless, improving the MDD diagnostic criteria may only be achievable through epistemic iterations^63^, a series of efforts to characterize specific genetic signals for the deepest available MDD definition and, in turn, refine our definition of MDD. Our efforts to improve GWAS power and specificity in noisy biobanks advances this process.

Our implementation of phenotype integration uses shallow MDD phenotypes to improve power for LifetimeMDD, and as such its specificity is limited by the specificity of LifetimeMDD. Future statistical methods could go further and actually improve the specificity of existing phenotypes. We have taken a complementary step in this direction by residualizing latent factors from SoftImpute, which revealed a specific locus from a shallow phenotype. This approach is akin to latent factor correction in genomic studies^22,68–71^ and it could be adapted to AutoComplete using, for example, Integrated Gradients^72^. However, it is challenging to remove non-specific signals without removing specific signals or, worse, introducing artificial signals due to collider bias^22,41,73^. This is especially true for disorders like MDD where epistemic uncertainty clouds what signals are most biomedically useful. A more principled approach is warranted.

Phenotype integration is broadly applicable to biobank-based genetic studies, which often evaluate a mixture of biomarker-, nurse-, GP-, specialist-, and/or patient-defined disorder statuses. Further, biobanks offer diverse disorder-relevant phenotypes, such as age of onset, medical procedures, prescriptions, environmental risk factors, family history, and socioeconomic measures. We expect phenotype integration to substantially improve GWAS power and PRS accuracy for many complex disorders. While the degree of specificity will vary between applications, this can be assessed with our novel PRS Pleiotropy metric. This metric characterizes a power-specificity tradeoff for MTAG, where adding more phenotypes generally increases power but sacrifices specificity. This empirical measure complements prior formal theoretical derivations of power-specificity tradeoffs in meta-analysis of heterogeneous traits^74^. In the future, PRS Pleiotropy can be used to evaluate newly constructed phenotypes, including manual combinations of existing measures^13^.

Importantly, our work has complex implications for equity in genetic studies and clinical care. On the one hand, EHR-derived phenotypes have a history of exacerbating inequities that continues to this day^75^. Moreover, phenotype integration uses a reference phenome, which has the potential to propagate systematic biases present in biobank data. On the other hand, we found that phenotype integration can improve PRS portability across ancestries. However, the results are less clear than those in European ancestries, highlighting the need for greater sample sizes in diverse ancestries and better methods for cross-ancestry portability. Additionally, mounting evidence suggests that portability can be improved by homing in on causal biology using transcriptomics^76^, genomic annotations^77^, or fine-mapping^78^. Therefore, careful extensions of our approach, such as residualizing phenome-wide factors, have the potential to improve portability by eliminating confounders. Given the extreme Euro-centric biases in available genomics data, these and other statistical approaches to improve the utility of PRS for all people are urgently needed.

## Methods

### Phenotypes used in phenotype imputation

We considered 217 relevant phenotypes to impute LifetimeMDD in 337,127 individuals of white British ancestry in UKB (**Supplementary Table 1**). These include: a) LifetimeMDD as defined in Cai et al 2020^10^; b) minimal phenotyping definitions of depression based on help-seeking, symptoms, self-reports, and/or electronic health records (EHR) as defined in Cai et al 2020^10^; c) individual lifetime and current MDD symptoms from the Composite International Diagnostic Interview – Short Form (CIDI-SF)^79^ and Patient Health Questionnaire (PHQ9) from which we derived LifetimeMDD; d) psychosocial factors; e) self-reported comorbidities; f) family history of common diseases; g) early life factors h) socioeconomic phenotypes; i) lifestyle and environment phenotypes; j) social support status; and k) demographic features including age, sex, UKBiobank assesement centre as a proxy for geographical residence, and 20 genetic PCs. These phenotypes are selected based on their established relevance to MDD, and are all collected through either the Touchscreen questionnaire completed at the assessment centre or through the online mental health follow-up questionnaire (MHQ). All UKBiobank data fields, sample sizes and prevalence of binary outcomes are detailed in **Supplementary Table 1**, and we report levels of missingness for all inputs for multi-phenotype imputation in **Extended Data Figure 1**. For PRS pleiotropy analyses, we excluded the 20 genetic PCs, 22 assessment centers, and genotyping array.

### Phenotype imputation with SoftImpute

We fit SoftImpute with the ALS method^23^ on the 217 phenotypes comprising the MDD-related phenome in UKB, using cross-validation to optimize the nuclear norm regularization parameter. We used our prior approach to make the cross-validation more realistic by copying real missingness patterns instead of completely random entries^16,80^, which provides far more realistic estimates of imputation accuracy (**Extended Data Figure 1**). We previously studied SoftImpute at a smaller scale in comprehensive simulations and several real datasets^16^, and we have since used it in several larger studies^16,80,81^. Overall, SoftImpute is extremely simple, robust, and scalable. We summarize the SoftImpute model fit by the latent factors (**Figure 3C**) and the variance they explain (**Figure 3A**), which are akin to the eigenvectors (or PCs) and eigenvalues of the phenotype covariance matrix, respectively. We also estimate the prediction strength (**Figure 3B**), which is the squared-correlation between two latent factors estimated after splitting the sample into two non-overlapping halves (*R^2^*). We define the effective sample size post-imputation as Nobs + Nmiss\**R^2^*, analogous to genotype imputation^16,17,24^; we note that this approximates the power-equivalent number of observed phenotypes.

### Phenotype imputation with AutoComplete

We developed a new deep-learning based method, AutoComplete, in a companion paper (An et al in submission). AutoComplete consists of several fully-connected layers with nonlinearities and learns to optimize reconstruction of realistically held-out missing entries. The model is fully differentiable and is fit using stochastic gradient descent. Unlike SoftImpute, Autocomplete’s objective function models binary phenotypes. As with SoftImpute, the hyperparameters for AutoComplete were determined through cross-validation on realistically held-out missing data. In this paper, we focus on its application to imputing LifetimeMDD.

### GWAS on observed or imputed phenotypes

GWAS on directly-phenotyped and imputed phenotypes in UKB was performed using imputed genotype data at 5,781,354 SNPs (minor allele frequency > 5%, INFO score > 0.9) using logistic regression and linear regression implemented in PLINK v2^82^ for binary and quantitative traits respectively. We used 20 PCs computed with flashPCA^83^ on 337,129 White-British individuals in UKB and genotyping arrays as covariates for all GWAS (see **Supplementary Note** for details in sample and genotype QC in UKB). To test for heterogeneity between genetic effects found in GWAS on observed LifetimeMDD and imputed measures of MDD from SoftImpute (Soft-ImpOnly) and AutoComplete (Auto-ImpOnly), we performed a random effect meta-analysis using METASOFT^32^ and tested for heterogeneity between effect sizes at each SNP.

### SNP-based heritability and genetic correlation

To test for SNP-based heritability of each phenotype and the genetic correlation between pairs of phenotypes, LD score regression implemented in LDSC v1.0.11^31,84^ was performed on the GWAS summary statistics using in-sample LD scores estimated in 10,000 random white British UKB individuals at SNPs with MAF > 5% as reference. For MTAG results, we used the effective sample size estimated in MTAG as sample size entry in LDSC; for all other GWAS, we use the actual sample size. When we estimate the liability-scale heritability, we assume the population prevalence of binary phenotypes equal their prevalence in UKB. For all GWAS, we have indicated their effective sample sizes accounting for imbalance between cases and controls (N_eff_ = 4/(1/N_cases_ + 1/N_controls_)) in **Supplementary Tables 5** and **6**. We note this is different from the imputation-related definition of effective sample size, and also differs from MTAG’s definition.

### In-sample PRS prediction of phenotypes in UKB with 10-fold cross validation

We performed SoftImpute^23^ and AutoComplete imputations 10 times, each time using 90% of the individuals in the input phenotype matrix, built PRS from GWAS results from this with PRSice v2^85^, and evaluated predictive accuracy for observed LifetimeMDD and the depression-related phenome (217 phenotypes, used as input in imputation) in the held-out 10%. For MTAG^18^, we performed GWAS on each set of input phenotypes (as shown in **Figure 4**) 10 times, each time using 90% of the individuals in UKB. We then ran MTAG on GWAS summary statistics in this 90%, built PRS from the resulting MTAG summary statistics with PRSice v2, and evaluated predictive accuracy for observed LifetimeMDD in the held-out 10%. For all PRS predictions, we used 20 genomic PCs and the genotyping array used as covariates. For binary phenotypes, including LifetimeMDD, we evaluated accuracy using Nagelkerke’s *R^2^*. For all quantitative phenotypes, including neuroticism, we evaluated accuracy using ordinary *R^2^*.

### PRS prediction of phenotypes in UKB from external GWAS summary statistics

We construct PRS from MDD GWAS summary statistics from PGC29^2^, iPSYCH^14^, and 23andMe^5^, as detailed in **Supplementary Table 6**, and predicted phenotypes in UKB using PRSice v2, using 20 genomic PCs and the genotyping array used in UKB as covariates. For each of these studies, we use only SNPs with imputation INFO score > 0.9 and MAF > 5% for constructing PRS. For binary phenotypes, including LifetimeMDD, we evaluated accuracy using Nagelkerke’s *R^2^*. For all quantitative phenotypes, including neuroticism, we evaluated accuracy using ordinary *R^2^*. We calculated PRS Pleiotropy on all secondary phenotypes (not including LifetimeMDD) used in imputation except for 20 PCs, array, assessment centers (in total 172 phenotypes).

## Extended Data Figures

**Extended Data Figure 1:**
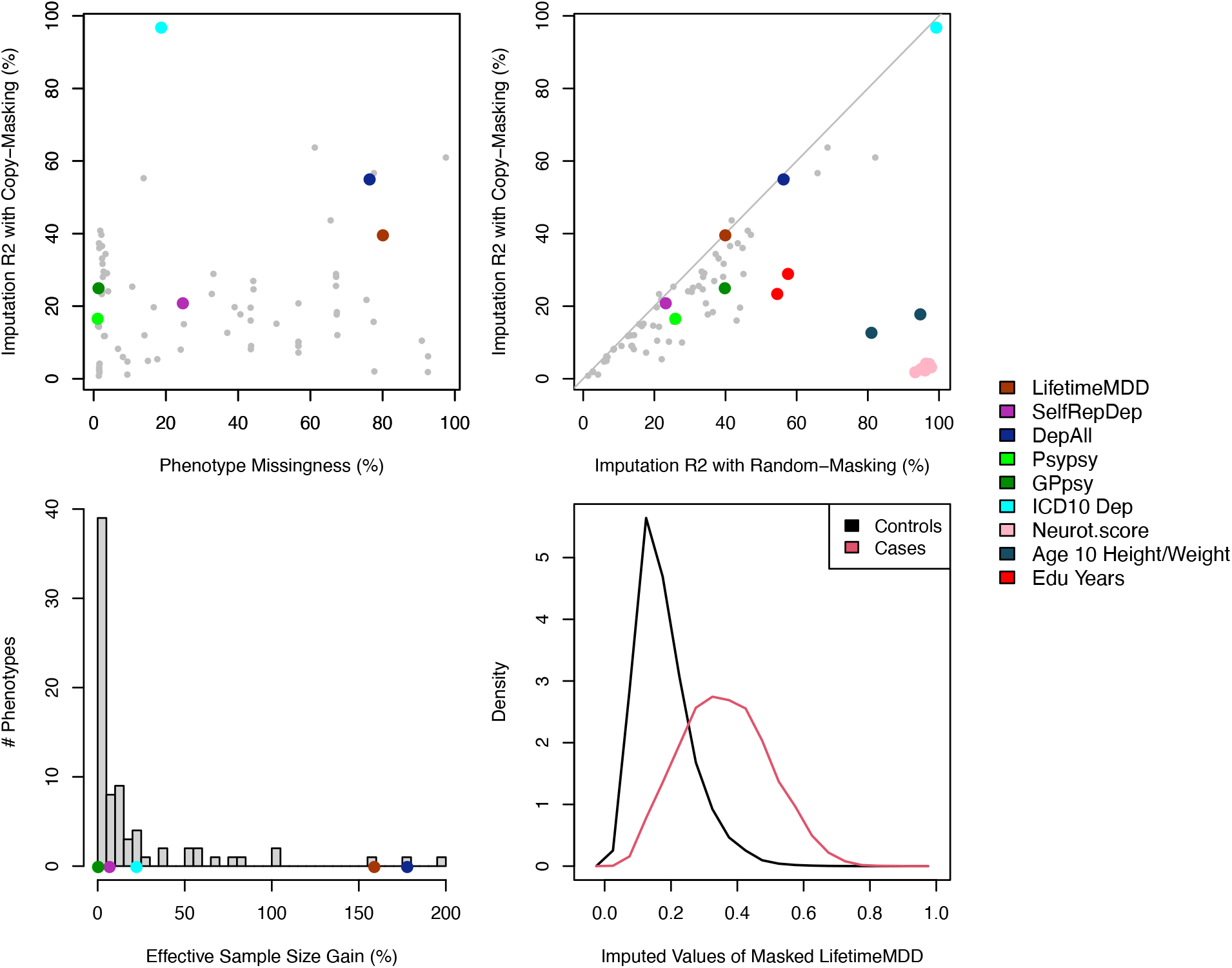
Imputation accuracy metrics across our depression-relevant UKB phenome. **(A)** Scatter plot of estimated imputation accuracy against phenotype missingness. **(B)** Scatter plot of estimated imputation accuracy using our copy-masking approach against naive estimates masking entries uniformly at random. **(C)** Distribution across phenotypes of gained effective sample size from phenotype imputation. (D) Distribution of imputed LifetimeMDD values for held-out observations, which informally reflect the probabilities of having LifetimeMDD.

**Extended Data Figure 2:**
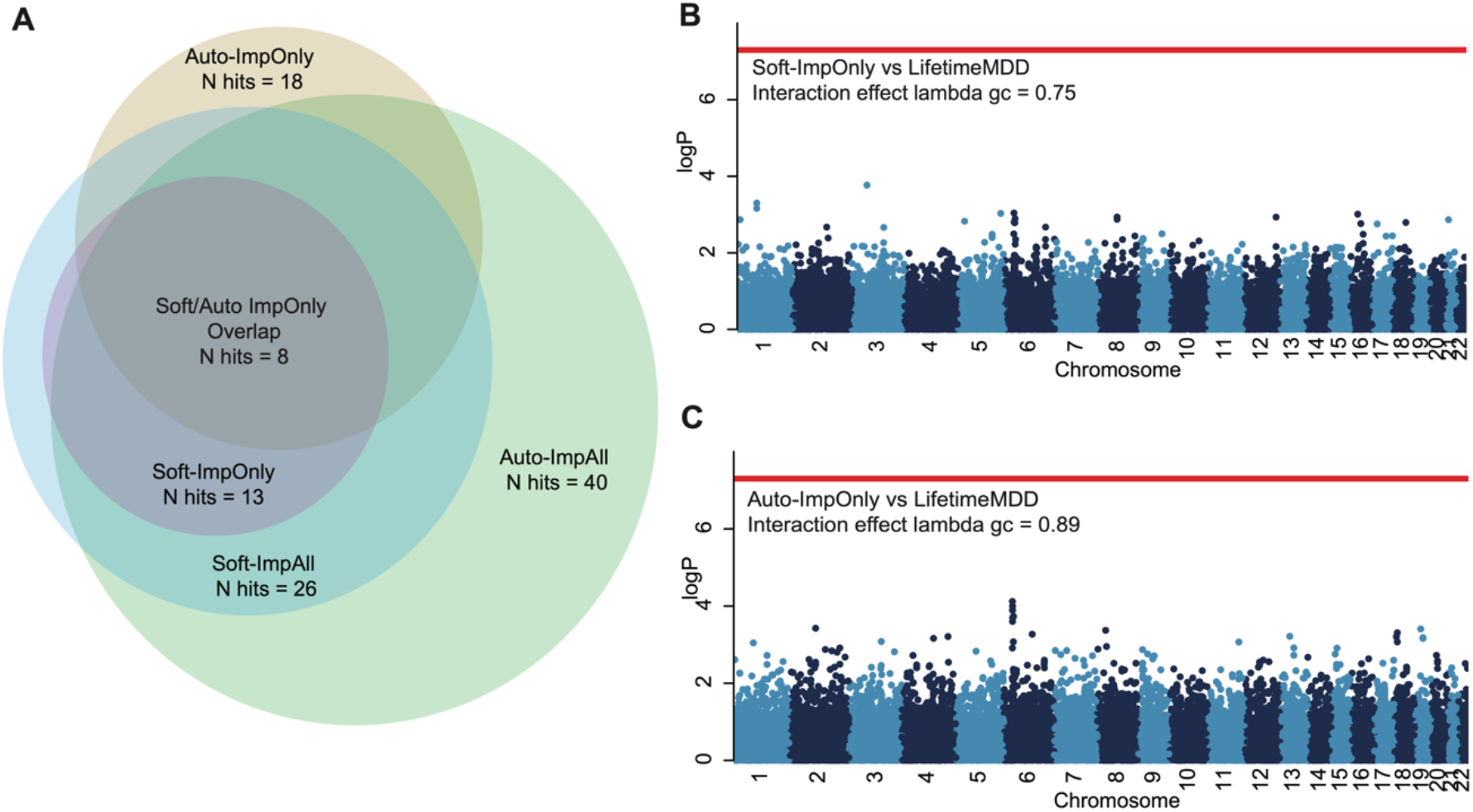
**(A)** Venn diagram showing the overlap of GWAS loci identified from GWAS on ImpOnly and ImpAll measures of LifetimeMDD from Softimpute and Autocomplete; **(B,C)** Manhattan plots of Cochran’s Q statistic P value for heterogeneity, obtained through a random effect meta-analysis performed with METASOFT, between genetic effects identified from GWAS on observed LifetimeMDD and GWAS on ImpOnly measures of LifetimeMDD from Softimpute or Autocomplete; red line shows the genome-wide significance threshold corresponding to P value 5×10^-8^.

**Extended Data Figure 3:**
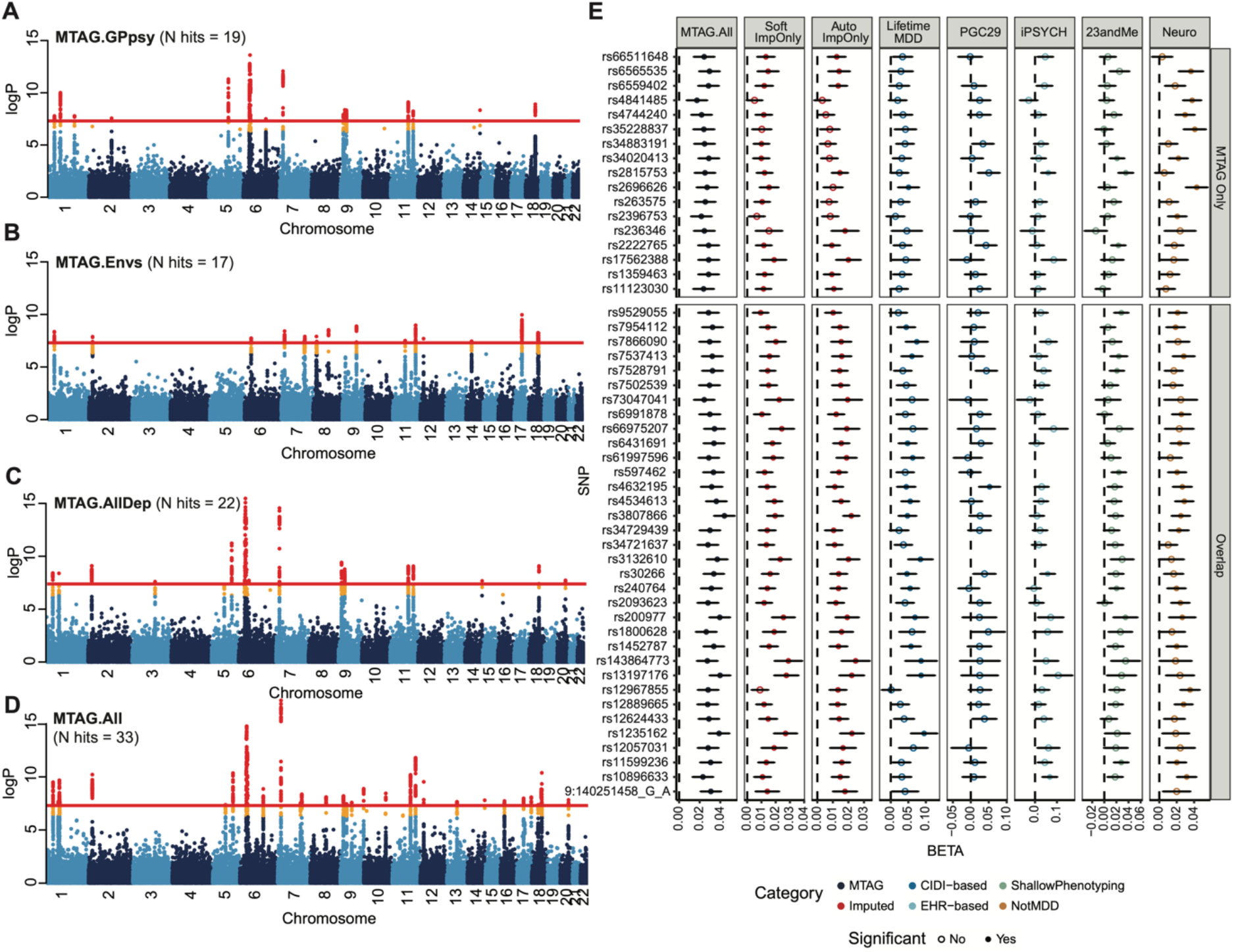
**(A-D)** Manhattan plots showing MTAG results for LifetimeMDD for the MTAG runs: MTAG.GPpsy, MTAG.Envs, MTAG.AllDep and MTAG.All, descriptions of which are shown in **Figure 4A**; red line shows the genome-wide significance threshold corresponding to P value 5×10^-8^; **(E)** Replication of GWAS effect sizes for LifetimeMDD for loci identified in MTAG runs only and those that overlap between MTAG and imputation (both Softimpute and Autocomplete), in observed LifetimeMDD and external MDD studies from PGC, iPSYCH, and 23andMe. All error bars indicate 95% confidence intervals.

**Extended Data Figure 4:**
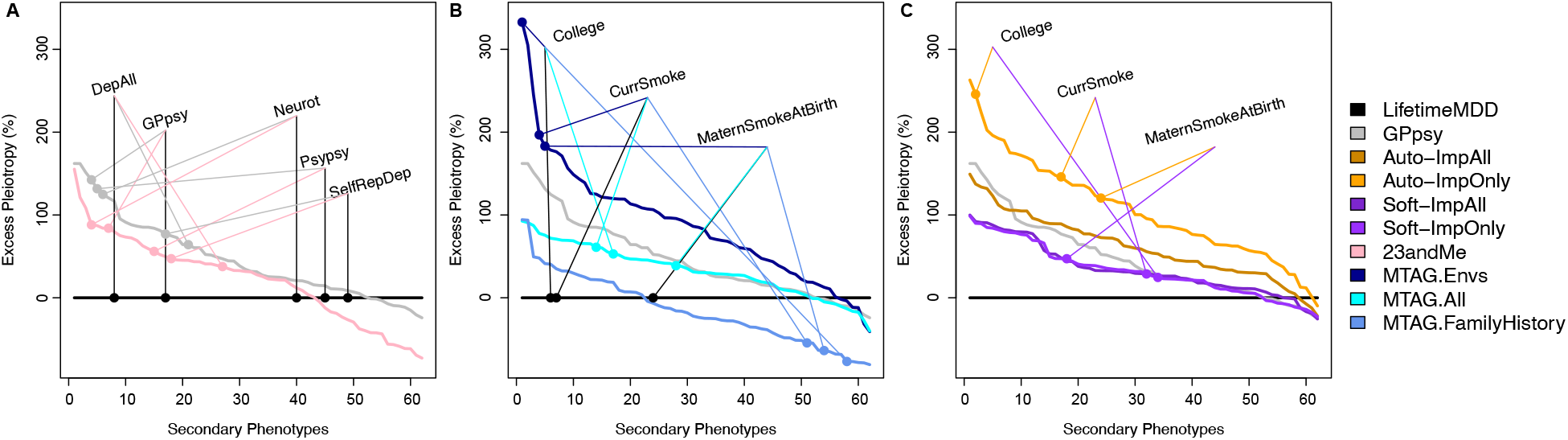
Excess PRS Pleiotropy of a PRS relative to the LifetimeMDD PRS. PRS Pleiotropy is defined as the PRS prediction ratio for a secondary trait relative to observed LifetimeMDD (PRS Pleiotropy:= *R^2^*_secondary_/*R^2^*_LifetimeMDD_), and excess pleiotropy is the increase in pleiotropy relative to the LifetimeMDD PRS (Excess PRS Pleiotropy:= (PRS Pleiotropy - LifetimeMDD PRS Pleiotropy)/LifetimeMDD PRS Pleiotropy). Plots are ordered by Excess PRS Pleiotropy for each phenotype PRS. **(A)** The PRS derived from GPpsy and 23andMe are less specific to LifetimeMDD than the LifetimeMDD PRS, especially for shallow MDD phenotypes and neuroticism **(B)** MTAG.Envs has high Excess PRS Pleiotropy to secondary traits like college education, smoking, and maternal smoking, while MTAG.FamilyHistory actually reduces PRS Pleiotropy for these traits. **(C)** Both ImpOnly and ImpAll SoftImpute phenotypes show lower Excess PRS Pleiotropy than GPpsy, while ImpAll GWAS from Autocomplete is comparable to GPpsy.

**Extended Data Figure 5:**
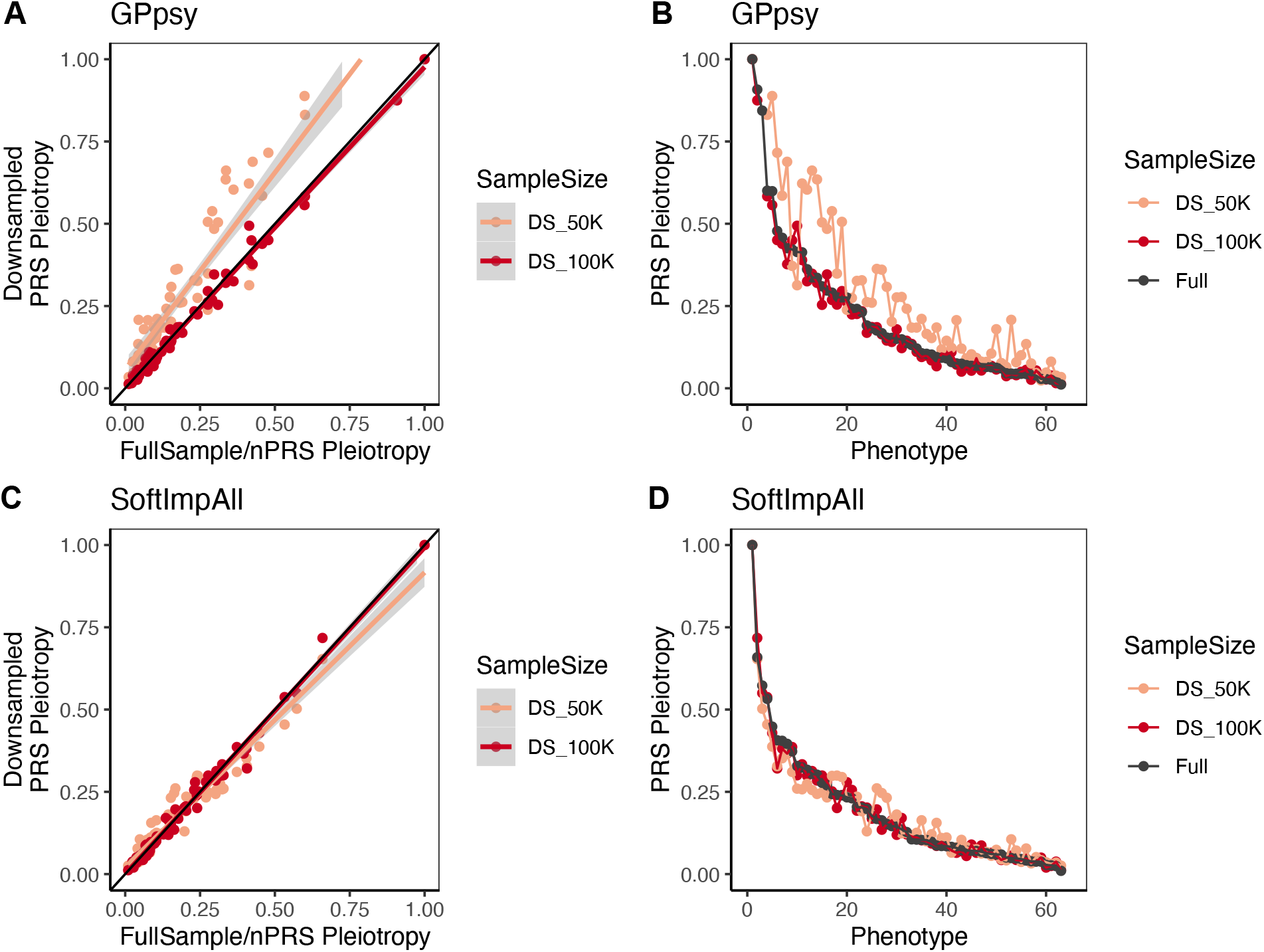
PRS Pleiotropy of **(A,B)** GPpsy (N=332,629) and **(C,D)** Soft-ImpAll (N=337,126) at full sample size and down-sampled to N=50K and 100K for 62 phenotypes in UKB. Plotted values are mean PRS Pleiotropy from 10-fold cross-validation in UKB.

**Extended Data Figure 6:**
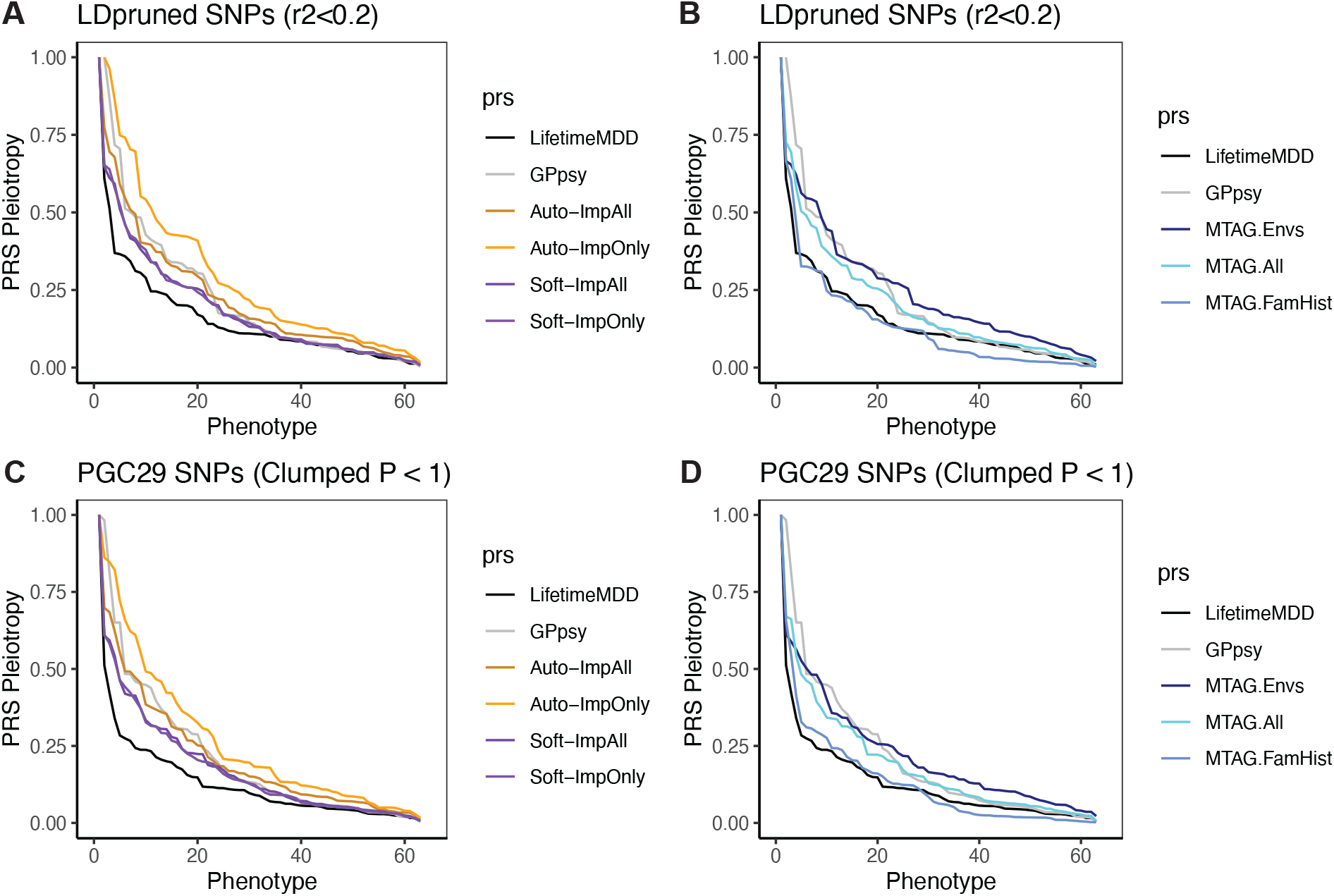
PRS Pleiotropy of imputed and MTAG GWAS for 62 phenotypes in UKB (which are significantly predicted by at least one full-sample PRS, as shown in Figure 6), where the PRS is constructed with **(A,B)** 136,563 LD-pruned SNPs (r2 < 0.2) in UKB, and **(C,D)** 91,315 SNPs from PGC29 GWAS clumped at P_threshold_ = 1; plotted values are mean PRS Pleiotropy from 10-fold cross-validation in UKB.

## Author contributions

AD and NC wrote the paper. AD, JF, KSK, and NC designed the study. MT performed ATLAS analyses, UA and SS performed AutoComplete imputation. AD and NC performed all other analyses. MK, VA, TW, and AJS supported iPSYCH analyses. All authors reviewed the paper.

## Acknowledgements

AD, SB, JF, AJS, SS, KSK, NC are supported by R01MH130581 from the NIH. MT is supported in part by the NIH Training Grant in Genomic Analysis and Interpretation T32HG002536. UA and SS are supported in part by III-1705121, CAREER-1943497, and R35GM125055. VA is supported by Lundbeck foundation postdoctoral grant R380-2021-1465. AJS is supported by Lundbeckfonden Fellowship R335-2019-2318. RB is supported by T32-NS048004 from the NIH. The iPSYCH team was supported by grants from the Lundbeck Foundation (R102-A9118, R155-2014-1724, and R248-2017-2003), NIMH (1R01MH124851-01) and the Universities and University Hospitals of Aarhus and Copenhagen. The Danish National Biobank resource was supported by the Novo Nordisk Foundation. High-performance computer capacity for handling and statistical analysis of iPSYCH data on the GenomeDK HPC facility was provided by the Center for Genomics and Personalized Medicine and the Centre for Integrative Sequencing, iSEQ, Aarhus University, Denmark. The authors gratefully acknowledge the support of all collaborators and participants in UKB, iPSYCH, CONVERGE and UCLA ATLAS who made this work possible.

## Declaration of interests

The authors report no financial relationships with commercial interests.

## Ethical approval

This research was conducted under the ethical approval from the UK Biobank Resource under application no. 28709 and 33217. The use of iPSYCH data follows standards of the Danish Scientific Ethics Committee, the Danish Health Data Authority, the Danish Data Protection Agency, and the Danish Neonatal Screening Biobank Steering Committee. Data access was via secure portals in accordance with Danish data protection guidelines set by the Danish Data Protection Agency, the Danish Health Data Authority, and Statistics Denmark. Retrospective data collection and analysis for ATLAS was approved by the UCLA IRB^47^. Patient Recruitment and Sample Collection for Precision Health Activities at UCLA is an approved study by the UCLA Institutional Review Board (UCLA IRB17-001013). All necessary patient/participant consent has been obtained and the appropriate institutional forms have been archived. The CONVERGE (China, Oxford, and VCU Experimental Research on Genetic Epidemiology) study was approved by the ethical review boards of Oxford University and participating hospitals. All participants provided written informed consent^7^.

## Data availability

UKBiobank genotype and phenotype data used in this study are from the full release (imputation version 2) of the UKBiobank Resource obtained under application no. 28709 and 33217. We used publicly available summary statistics from PGC29 and 23andMe from the Psychiatric Genomics Consortium (https://www.med.unc.edu/pgc/results-and-downloads), as well as summary statistics for affective disorders in the iPSYCH2012 cohort (https://doi.org/10.6084/m9.figshare.20517330), with references in **Supplementary Table 4**. The individual-level CONVERGE, Danish and UCLA datasets are not publicly available due to institutional restrictions on data sharing and privacy concerns. Summary statistics of all GWAS used in this paper are available at: https://doi.org/10.6084/m9.figshare.19604335.

## Code availability

Publicly available tools that are used in data analyses are described wherever relevant in Methods and Reporting Summary. Custom code for softImpute imputation of the MDD-relevant phenome and calculating PRS Pleiotropy are available at https://github.com/andywdahl/mdd-impute and https://github.com/caina89/MDDImpute. The Autocomplete software is available at: https://github.com/sriramlab/AutoComplete.

## Notes

### Competing Interest Statement

The authors have declared no competing interest.

### Summary of Updates

1. We evaluated three alternatives to our phenotype imputation matrix: (1) sex-stratified; (2) adding BMI; and (3) restricting to MTAG phenotypes (Supplementary Figure 2). (1) slightly hurt performance, (2) had little impact, and (3) performed far worse. 2. We significantly improved our presentation of the MTAG results, especially by expanding on our description of its utility in the Discussion and by modifying section titles. 3. We now publicly release our GWAS summary statistics. 4. We clarified/corrected our use of a few key terms, e.g. effective sample size and phenotype integration. 5. We thoroughly evaluated the variances and correlations of the imputed phenotypes (Supplementary Figure 1). 6. We added a major caveat to our Discussion on the potential for phenotype imputation to bias downstream analyses that others might perform, as well as potential solutions for these future directions. 7. We formally show that the increase in GWAS power from phenotype integration is statistically significant (Supplementary Figure 3). 8. We expanded our Discussion and added relevant references to place our work in context of prior efforts to improve power for MDD GWAS, including proxy GWAS (GWAX) and combining endorsements of multiple depression measures. 9. We extensively characterized PRS Pleiotropy as a function of sample size (Extended Data Figure 5). Most importantly, our results confirm that the conclusions in our paper are robust to sample size differences between our PRS. More broadly, our results characterize the complex interplay between sample size, p-value threshold, and the number of SNPs in a PRS (Supplementary Figures 10-11, Extended Data Figure 6). 10. We added two key references to prior work on MTAG; a theoretical result that complements our empirical results on power-specificity tradeoffs, and a smaller-scale observation consistent with our result that MTAG inflates genetic correlation. 11. We corrected a small but important overstatement in our previous submission on the relationship between portability and biological causality.

