## SupplementaryMaterials for "Phenotype integration improves power and preserves specificity in biobank-based genetic studies of MDD"

**Supplementary Material**

**Supplementary Note** 3

Imputed genotype quality control in UKBiobank 3

Sample filtering in UKBiobank 3

Tuning parameters in SoftImpute 3

Selection of phenotypes for MTAG runs 4

PRS prediction of MDD in non-White British individuals in UKBiobank 4

PRS prediction of MDD in external cohorts 5

*iPSYCH* *5*

*UCLA ATLAS* *6*

*CONVERGE* *7*

Impact of sample size on PRS Pleiotropy 8

Impact of P value threshold on PRS Pleiotropy 8

**Supplementary Figures** 10

Supplementary Figure 1. 10

Supplementary Figure 2. 11

Supplementary Figure 3. 12

Supplementary Figure 4 13

Supplementary Figure 5 14

Supplementary Figure 6 15

Supplementary Figure 7 16

Supplementary Figure 8 17

Supplementary Figure 9 18

Supplementary Figure 10 19

Supplementary Figure 11 20

**Supplementary Tables** 21

**Supplementary References** 22

### Supplementary Note

#### Imputed genotype quality control in UKBiobank

We performed stringent filtering on imputed variants (version 3) in the UKBiobank[1] used for GWAS in this study. We hard-called genotypes from imputed dosages at 9,720,240 biallelic SNPs with imputation INFO score greater than 0.9, MAF greater than 0.1%, and P value for violation of Hardy-Weinberg equilibrium > $10^{-6}$, with a genotype probability threshold of 0.9 (anything below would be considered missing). Of these, 5,776,313 SNPs are common (MAF > 5%). We consistently use these SNPs for all analyses in this study.

#### Sample filtering in UKBiobank

Of all 502,637 samples in UKBiobank full release[1], we performed the following QC steps to select the samples for use in our analyses. We first removed samples that were not included in the UKBiobank full release PCA analysis, which includes samples that were indicated as "het.missing.outliers" ("Indicates samples identified as outliers in heterozygosity and missing rates, which indicates poor-quality genotypes for these samples"), "excess.relatives" ("Indicates samples which have more than 10 putative third-degree relatives in the kinship table"), and whose "Submitted.Gender" were different from "Inferred.Gender". Applying these filters brought the sample size down to 407,219. We checked that the remaining sample contains only one out of any pair or group of related individuals with relatedness > 0.05. We then selected individuals indicated to be "in.white.British.ancestry.subset" ("Indicates samples who self-reported 'White British' and have very similar genetic ancestry based on a principal components analysis of the genotypes"), resulting in a sample size of 337,545. We then removed 337 individuals indicated as having "putative.sex.chromosome.aneuploidy" ("Indicates samples identified as putatively carrying sex chromosome configurations that are not either XX or XY")[1]. Finally, we removed 418 individuals who have withdrawn their consent for use of their genetic data in analyses, arriving at our final set of 337,127 individuals passing QC. Of these individuals, 37,035 were part of UK Biobank Lung Exome Variant Evaluation (UKBiLEVE)[2], a study for chronic obstructive pulmonary disease (COPD). We retain all samples in UKBiLEVE, but as they are genotyped using a custom array optimized for coverage over regions implicated in lung health and disease[2], we consistently use the genotyping array as a covariate in all our analyses.

#### Tuning parameters in SoftImpute

SoftImpute[3] operates by finding an optimal low-rank approximation, X, to the partially observed phenotype matrix, Y, by solving $||Y_o - X_o||_2^2 + \lambda||X||_*$ where the o subscript indicates observed phenotype entries in Y, λ is a regularization parameter, $||\cdot||_2$ indicates the $l_2$ norm, and $||\cdot||_*$ indicates the nuclear norm of a matrix, i.e., the sum of its singular values. The nuclear norm is a computationally and statistically efficient alternative to a non-convex rank constraint on X, as is common in PCA. SoftImpute is the matrix version of lasso regression, wherein an $l_1$ penalty is used to relax a non-convex constraint on the number of active regression features. We tune λ to maximize imputation accuracy on held-out test data, which is realistically sampled to ensure representative patterns of missingness[4].

**Selection of phenotypes for MTAG runs**

We did not include all 216 phenotypes used in imputation in MTAG analyses, because it was impractical to perform individual GWAS on all 216 phenotypes. Instead, we wanted to test for how agnostic MTAG is for input phenotypes, and therefore selected 6 different sets of input phenotypes for MTAG, as shown in **Figure 4**. In these 6 sets of input phenotypes we tried to capture the different axes of shared genetic contributions between input phenotypes and MDD, namely: a) the sharing between deep and shallow phenotyping definitions of MDD (including GPpsy, and all other definitions of depression as previously specified in Cai et al 2020[5], in MTAG.GPpsy and MTAG.AllDep); b) the sharing between observed LifetimeMDD and family history of MDD (including severe depression in mother, father or siblings, in MTAG.FamilyHistory); and c) the sharing between environmental factors and personality traits and MDD (including neuroticism score, townsend deprivation index, recent stressful life events and lifetime trauma as specified in Cai et al 2020[5], in MTAG.Envs); and d) the combinations of the previous categories (in MTAG.AllDep+Envs and MTAG.All). Definitions of each of the phenotypes in the 6 sets are detailed in **Supplementary Table 1**. We recognise that we do not provide a comprehensive list of phenotypes one can potentially use in MTAG to improve GWAS power and PRS prediction accuracy for MDD. Rather, here we focus on investigating the effects of both the number and nature of MTAG input on the results.

**PRS prediction of MDD in non-White British individuals in UKBiobank**

We identified 150,213 individuals who did not self-identify as White British in the UKBiobank[1]. In these individuals, we removed from their imputed genotypes regions of the genome that contained previously identified structural variants[6], then hard-called genotypes from imputed dosages at 5,327,974 biallelic SNPs with imputation INFO score greater than 0.9, MAF greater than 0.1%, and P value for violation of Hardy-Weinberg equilibrium > 10-6, with a genotype probability threshold of 0.9 (anything below would be considered missing).

To identify participants of European (EUR), African (AFR) and East Asian (ASN) ancestries, we selected 70,718 LD-pruned common SNPs (MAF > 5%, P value for HWE > $10^{-6}$, LD r2 < 0.1) the autosomes that overlap with SNPs in the 1000 Genomes Project Phase 3 (1000G)[7] from 150,213 non-White British individuals in UKBiobank, then projected all individuals onto PCs obtained from 1000G samples using loadings at these SNPs calculated with LDAK v5[8] (**Supplementary Figure 5**). We identified 122,710, 8,040 and 2,530 individuals who clustered with the individuals in 1000G with EUR, AFR and ASN ancestries respectively (**Supplementary Figure 5**). Of these individuals with non-White British ancestries, we removed 652 individuals indicated as having "putative.sex.chromosome.aneuploidy" ("Indicates samples identified as putatively carrying sex chromosome configurations that are not either XX or XY")[1], as well as 71,143, 542 and 70 individuals with EUR, AFR and ASN ancestries respectively with relatedness greater than 0.05 ('Kinship' score in UKBiobank full release sample QC file) with any of the 337,127 individuals with White-British ancestries in UKbiobank we use in our main analyses, or to each other. We therefore retain 51,567, 7,497 and 2,460 unrelated individuals of EUR, AFR and ASN ancestries for our analyses. For each group, we performed PCA with all 5,327,974 biallelic SNPs using flashPCA[9] (**Supplementary Figure 5**), and showed that they contain few individuals of discordant self-reported ancestries. We use the top 20 PCs from each group as covariates for all following analyses.

Of these individuals, a subset has answered the qualifying questions in the online mental health questionnaire for deriving a LifetimeMDD disease status[5]: for the EUR individuals, 10,193 individuals has LifetimeMDD disease status (Ncases = 2741, Ncontrols = 7452, prevalence = 0.27); for the ASN individuals, 334 individuals has a LifetimeMDD disease status (Ncases = 62, Ncontrols = 272, prevalence = 0.19); for the AFR individuals, 687 has a LifetimeMDD disease status, (Ncases = 122, Ncontrols = 565, prevalence = 0.18). For PRS analysis, we used 2,441,319, 1,808,453 and 2,025, 285 SNPs (MAF >= 0.05, INFO score >= 0.9, P value for HWE violation > 10-6) in the EUR, ASN and AFR individuals in UKBiobank respectively, and calculated PRS for each of the 15 summary statistics (phenotyped, imputed and MTAG GWAS) using PRSice v2[10], using the options --clump-kb 250kb --clump-p 1 --clump-r2 0.1 --interval 5e-05 --lower 5e-08. We used the top 20 genomic PCs from the EUR, ASN and AFR individuals in UKBiobank as covariates to control for population structure in each of the cohorts.

**PRS prediction of MDD in external cohorts**

***iPSYCH***

We first evaluated the transferability of PRS derived from phenotyped, imputed and MTAG summary statistics using two independent cohorts from the Integrative Psychiatric Research Consortium (iPSYCH) cohort (2012 and 2015i).

The Lundbeck Foundation initiative for Integrative Psychiatric Research (iPSYCH)[11,12] is a case-cohort study of all singleton births between 1981 and 2008 to mothers legally residing in Denmark and who were alive and residing in Denmark on their first birthday (N=1,657,449). The iPSYCH 2015 case-cohort comprises two enrollments from this base population. The iPSYCH 2012 case-cohort enrolled 86,189 individuals (30,000 random population controls; 57,377 psychiatric cases)[11]. The iPSYCH 2015i case-cohort expanded enrollment by an additional 56,233 individuals (19,982 random population controls; 36,741 psychiatric cases)[11,12]. DNA was extracted from dried blood spots stored in the Danish Neonatal Screening Biobank[13] and genotyping was performed on the Infinium PsychChip v1.0 array (2012) or the Global Screening Array v2 (2015i). Psychiatric diagnoses were obtained from the Danish Psychiatric Central Research Register (PCR)[14] and the Danish National Patient Register (DNPR)[15]. Diagnoses in these registers are made by licensed psychiatrists during in- or out- patient specialty care but diagnoses or treatments assigned in primary care are not included. Linkage across population registers, to parents where known, and to the neonatal biobank is possible via unique citizen identifiers of the Danish Civil Registration

System[16]. The use of this data follows standards of the Danish Scientific Ethics Committee, the Danish Health Data Authority, the Danish Data Protection Agency, and the Danish Neonatal Screening Biobank Steering Committee. Data access was via secure portals in accordance with Danish data protection guidelines set by the Danish Data Protection Agency, the Danish Health Data Authority, and Statistics Denmark. For this study, we use an unrelated, homogeneous ancestry subset of these data from the 2012 and 2015i cohorts. We include individuals with available genotypes, that passed our quality control, and were either a random control or diagnosed with major depressive disorder (MDD) as of Dec 31, 2015 (2012: Ncontrols = 23,371, Ncases = 18,879; 2015i: Ncontrols = 15,163, Ncases = 8,188; Total: Ncontrols = 38,534, Ncases = 27,067).

Genotype phasing, imputation, and quality control were performed in parallel in the 2012 and 2015i cohorts according to custom, mirrored protocols. Briefly, phasing and imputation were conducted using BEAGLEv5.1[17,18], both steps including reference haplotypes from the Haplotype Reference Consortium v1.1 (HRC)[19]. Quality control was applied prior to and following imputation to correct for missing data across SNPs and individuals, SNPs showing deviations from Hardy-Weinberg equilibrium in cases or controls, abnormal heterozygosity of SNPs and samples, genotype-phenotype sex dicordance, minor allele frequency (MAF), batch artifacts, and imputation quality. Kinship was detected within and across 2012 and 2015i cohorts using KING[20], censoring to ensure no second degree or higher relatives remained. Ancestry was examined using the smartpca module of EIGENSOFT[21], and PCA outliers from the set of iPSYCH individuals with both grandparents and four grandparents born in Denmark were excluded.

Using 5,210,642 and 5,222,714 SNPs (MAF >= 0.05, INFO score >= 0.9, P value for HWE violation > $10^{-6}$) on 42,250 individuals in iPSYCH2012 and 23,351 individuals in iPSYCH 2015i respectively, we calculated PRS for each of the 15 summary statistics (phenotyped, imputed and MTAG GWAS) using PRSice v2[10], using the options --clump-kb 250kb --clump-p 1 --clump-r2 0.1 --interval 5e-05 --lower 5e-08. We used the top 10 genomic PCs from all 42,250 and 23,351 individuals in iPSYCH2012 and iPSYCH2015i respectively as covariates to control for population structure in each of the cohorts.

Note that this analysis uses more recent releases of iPSYCH data than available in the summary statistics from Schork et al 2019[22], which we use for heritability and genetic correlation shown in **Figure 4**, and PRS analysis with published external summary statistics shown in **Figure 5A**.

#### *UCLA ATLAS*

We also evaluated the transferability of PRS on a diverse subset (n=19,657) of the individuals in the UCLA ATLAS cohort[23,24]. All individuals in the ATLAS cohort are genotyped at 673,148 genetic variants on a custom Illumina Global Screening (GSA) array that included a standard GWAS backbone and an additional set of pathogenic variants selected from ClinVar[23]; genotype imputation was performed on the Michigan Imputation Server[25], which performs phasing using Eagle v2.4[26] and performs imputation using the TOPMed Freeze5 imputation panel[27] and minimac4[28].

Individuals in the ATLAS dataset reported self-identified race/ethnicity (SIRE). We applied filtering on discrepancies between SIRE and clusterings of individuals with their first two genetic PCs (**Supplementary Figure 6**). Further, we removed individuals and variants with more than 1% missing values as well as variants that were not under Hardy-Weinberg equilibrium (p-value threshold $10^{-6}$). Ultimately, we included 1,997 unrelated samples and 5,770,863 variants for Asian-identifying individuals, 1,125 unrelated samples and 7,048,125 variants for Black-identifying individuals, 2,169 unrelated samples and 7,477,693 variants for Latino-identifying individuals, and 14,366 unrelated samples and 7,269,267 variants for White-identifying individuals in our analyses. The number of individuals with each Phecode in each SIRE category that are identified as cases or controls for ATLAS.DPR or ATLAS.MDD are shown in **Supplementary Table 7**.

As the UCLA ATLAS is derived from EHR, where disease status of participants are in the form of billing, insurance or ICD10 codes, many of the codes are repetitive or redundant. We applied a mapping between all the available billing, insurance and ICD10 codes to a standardized Phecode in ATLAS to represent the total number of individuals having a certain disease. For MDD disease status in the ATLAS EHR, we identified two Phecodes: Phecode 296.2 (ATLAS.DPR) for the superset of depressive illness, corresponding to ICD10 codes for depressive disorders as shown in **Supplementary Table 8**; and Phecode 296.22 (ATLAS.MDD) for the more stringently defined MDD disease status that corresponds to ICD10 codes for Major Depressive Disorder as shown in **Supplementary Table 8**. Of the 1,705 MDD cases in ATLAS.MDD, 1,690 are also cases in ATLAS.DPR (N=3,120).

To predict ATLAS.MDD or ATLAS.DPR in each of the SIRE categories, we obtained PRS from the individuals in ATLAS from each of the 15 summary statistics (phenotyped, imputed and MTAG GWAS) using PRSice v2[10], using the options --clump-kb 250kb --clump-p 1 --clump-r2 0.1 --interval 5e-05 --lower 5e-08. We used the top 10 genomic PCs from individuals in each of the SIRE categories individuals in ATLAS as covariates to control for population structure in each of the cohorts.

***CONVERGE***

Patients with recurrent MDD were recruited from 58 provincial mental health centers and psychiatric departments of medical hospitals in 45 cities and 23 provinces of China. Control subjects were recruited from multiple locations, including general hospitals and local community centers; all were screened and did not meet criteria for MDD, schizophrenia, or bipolar illness. Study participants were Han Chinese women with four Han grandparents. Case subjects were ages 30–60 and had at least two episodes of major depression meeting DSM-IV criteria, with the first episode between ages 14 and 50. The study was approved by the ethical review boards of Oxford University and participating hospitals. All participants provided written informed consent. Details on sample collection, phenotypes, and sequencing have been reported previously[29,30].

Using 4,570,966 SNPs (MAF >= 0.05, INFO score >= 0.9, P value for HWE violation > $10^{-6}$) on

10,502 individuals in CONVERGE (Ncases = 5,282, Ncontrols = 5,220, estimated population prevalence = 0.08), we calculated PRS for each of the 15 summary statistics (phenotyped, imputed and MTAG GWAS) using PRSice v2[10], using the options --clump-kb 250kb --clump-p 1 --clump-r2 0.1 --interval 5e-05 --lower 5e-08, and the top 10 genomic PCs from all 10502 individuals in CONVERGE as covariates to control for population structure.

**Impact of sample size on PRS Pleiotropy**

We directly investigated the effect of sample size by down-sampling. Specifically, we repeated our PRS pleiotropy analyses after down-sampling Soft-ImpAll (N=337,126) and GPpsy (N=332,629) to 50K and 100K. This involved 10-fold cross-validated GWAS and PRS construction, including tuning the optimal p-value threshold per each cross-validated PRS). We studied the same 62 phenotypes where all full-sampled PRS gave significant predictions (used in **Figure 6**)

We found that down-sampled PRS Pleiotropy are very highly correlated to full-sample PRS Pleiotropy (Soft-ImpAll: Pearson r for 50K = 0.98; Pearson r for 100K = 0.99; GPpsy: Pearson r for 50K = 0.95; Pearson r for 100K = 0.99). While GPpsy at N=50K had significantly higher mean PRS Pleiotropy (two-sided paired t-test P = $3.16 \times 10^{-11}$, mean difference = 10.2%), this difference vanishes at N=100K (P = 0.727, mean difference = 0.14%). Down-sampled Soft-ImpAll does not show significant difference in mean PRS Pleiotropy at either 50K (P = 0.90, mean difference = -0.06%) or 100K (P = 0.70, mean difference = 0.10%).

We show in **Extended Data Figure 5** the full and down-sampled PRS Pleiotropy across all 62 phenotypes ordered by their respective full-sample PRS Pleiotropy. Consistent with the above statistics, there were nontrivial fluctuations for N=50K yet surprisingly little fluctuation for N=100K. The variations were more extreme for GPpsy than Soft-ImpAll, which we attribute to differences in R2 for LifetimeMDD (which is likely the factor through which sample size acts).

Overall, we conclude that PRS Pleiotropy is qualitatively stable for a single PRS as a function of sample size (because correlations are high) and that it can be compared across different PRS for N >100K (though we cannot evaluate N>300K). In the context of our study, this is not problematic because only observed LifetimeMDD has N<100K (N=67K), which only means that our baseline is conservative.

**Impact of P value threshold on PRS Pleiotropy**

We then investigated how sample sizes would affect P value thresholds used for construction of PRS, and how that could affect PRS Pleiotropy measures.

Specifically, we investigated how sample size affects optimal P value thresholds, the number of included SNPs, and the variability of these statistics across folds of the data (**Supplementary Figures 10,11**). We find the following. First, the down-sampled PRS thresholds are more liberal and more variable (panels **A,B**). Second, the down-sampled PRS include more SNPs (panel **C,D**), which shows the net effect of more liberal thresholds (increasing number of SNPs used in PRS construction) and lower power (decreasing number of SNPs used in PRS construction). Third, down-sampled PRS gives more variable prediction $R^2$ across folds (panel **E**, as measured by coefficient of variation SD($R^2$/mean($R^2$)), which is expected but distinct from the fact that the $R^2$ decrease. Fourth, this results in higher variance in PRS Pleiotropy when using down-sampled PRS (panel **F**, especially for N=50K), which is expected but distinct from the fact that the average PRS Pleiotropy increases (especially for N=50K).

### Supplementary Figures

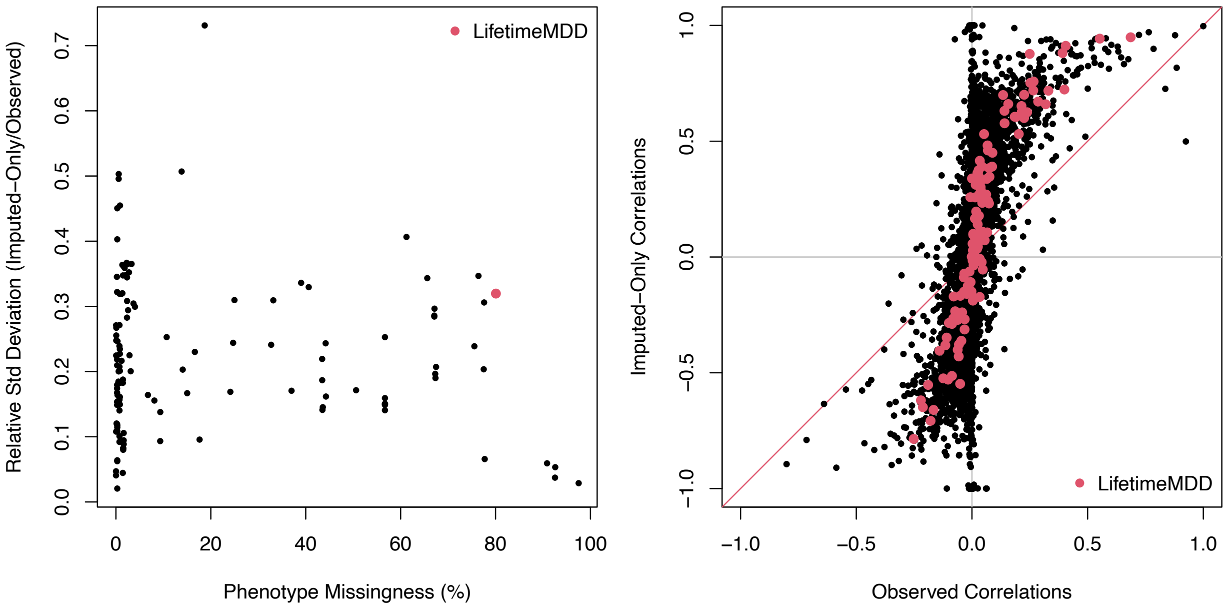


**Supplementary Figure 1.**

**(A)** Variance of SoftImpute imputed vs observed phenotypes. The variances were deflated by imputation, as expected. The average deflation in standard deviation was 20.3% (10th percentile=8.2%, 90th percentile=36.0%). **(B)** Correlation within imputed phenotypes vs within observed phenotypes. The correlations were inflated by imputation, as expected. The average 3.95 (10th percentile=1.35, 90th percentile=13.3). Note: the imputed covariances are actually smaller than the observed, but this is because their variances are smaller.

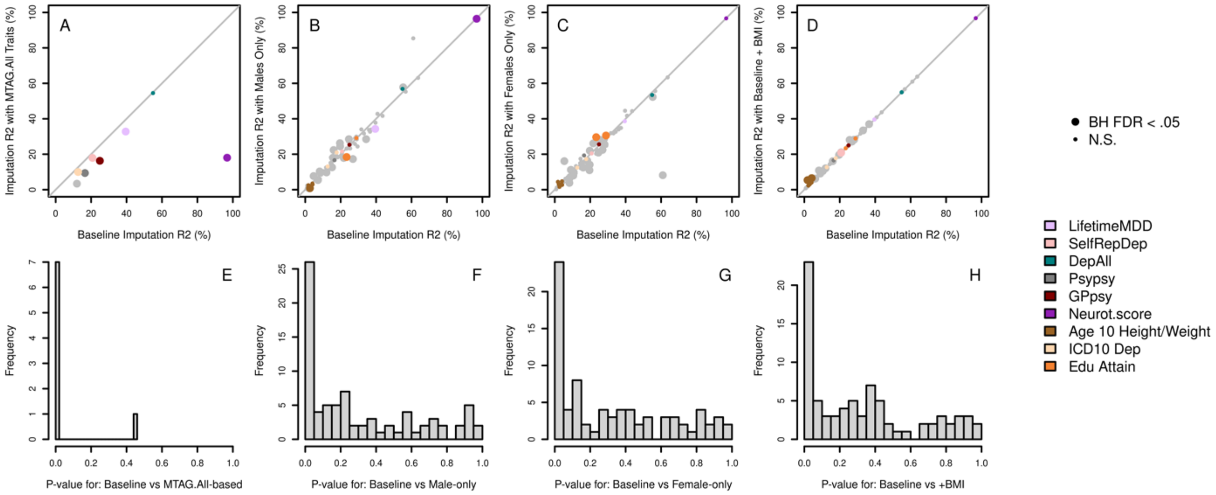


**Supplementary Figure 2.**

Comparison of our baseline imputation accuracy with softImpute to alternative approaches using different input phenotype matrices. Accuracy is compared to approaches that impute a phenotype matrix including: **(A)** only the MTAG.All traits plus demographic traits (age, sex, and 20 PCs); **(B)** only males; **(C)** only females; **(D)** adding a column for BMI. **(E-H)** show the P values comparing the baseline $R^2$ with the relevant alternative imputation approach. P values are calculated based on pooled t-tests across 10 replicates of copy-masked data with 1% added missingness (Methods) for **E-G**; **H** uses paired t-tests instead because the copy-masks are identical in both imputation approaches.

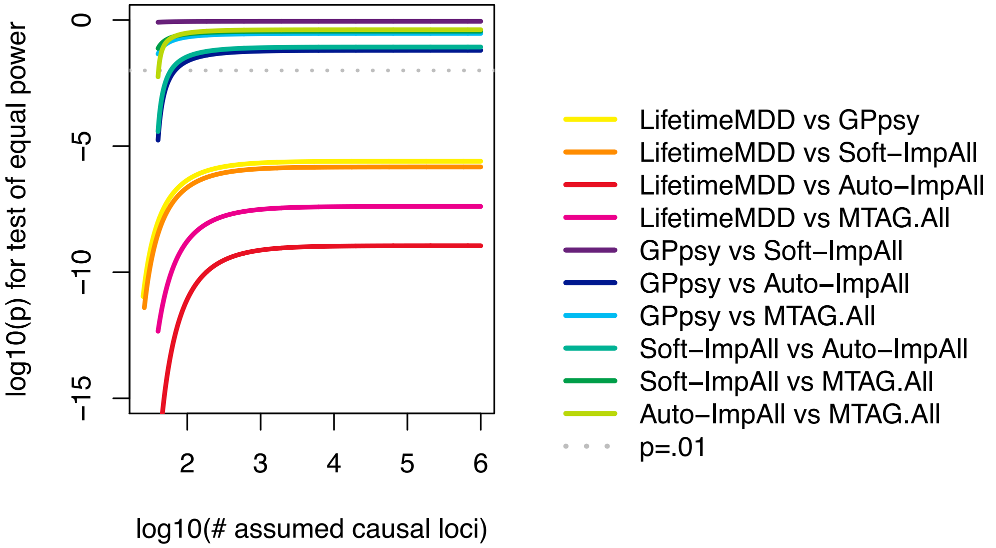


**Supplementary Figure 3.**

Binomial tests for equal power between each pair of 5 MDD GWAS in UKB on unrelated white British individuals. Each point on a curve tests the hypothesis that GWAS 1 and GWAS 2 have equal power to detect each of N causal loci. Specifically, for each number of causal loci N, we assume that the number of GWAS hits for each GWAS is independently distributed $n_i$~binomial(N,$p_i$) and then test whether n1=n2 using Pearson's chi-squared test. Our test shows that GPpsy and phenotype integration (imputed or MTAG.All) GWAS are all more powerful than the (observed) LifetimeMDD GWAS; except for implausibly few causal loci (e.g., N<100), our test does not detect different differences between phenotype integration GWAS.

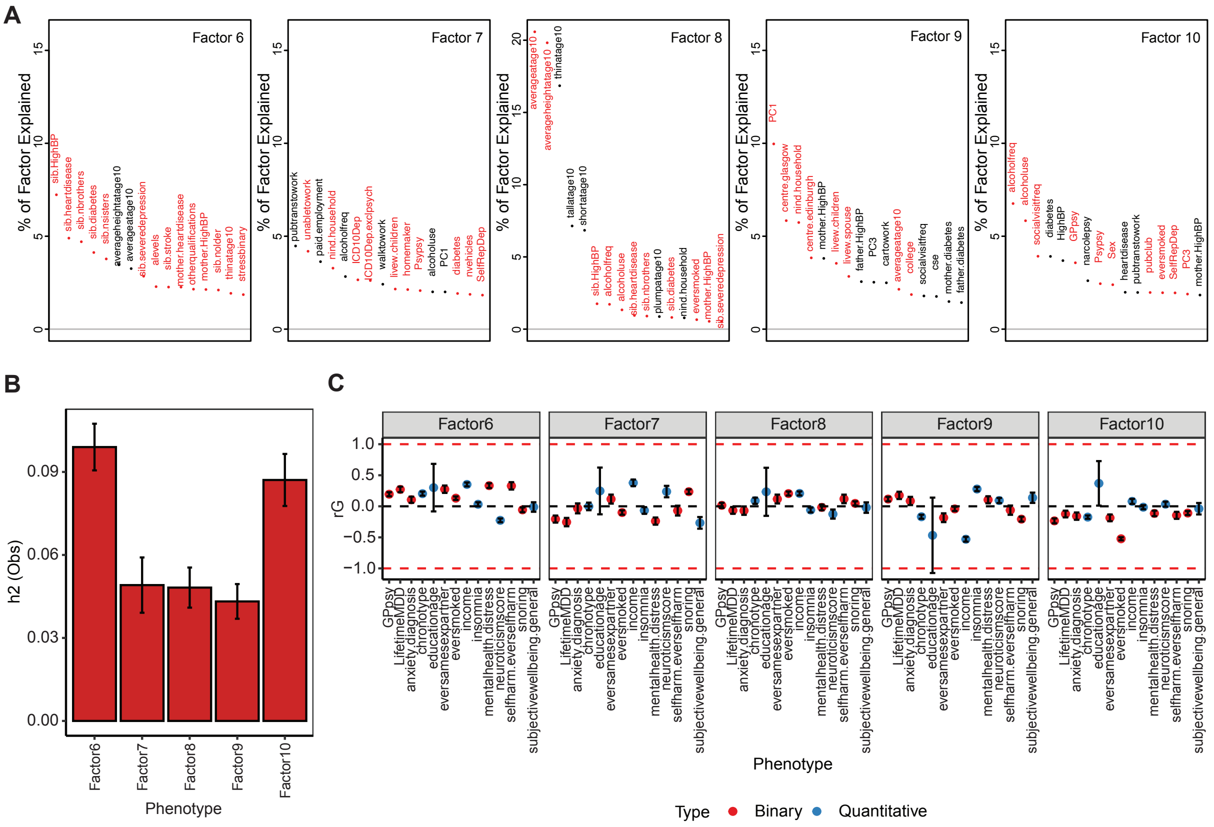

### Supplementary Figure 4

**(A)** Top phenotype loadings for the top 6-10 Softmpute factors; **(B)** Estimates of heritability and **(C)** genetic correlations of the top 6-10 SoftImpute factors to MDD-relevant traits.

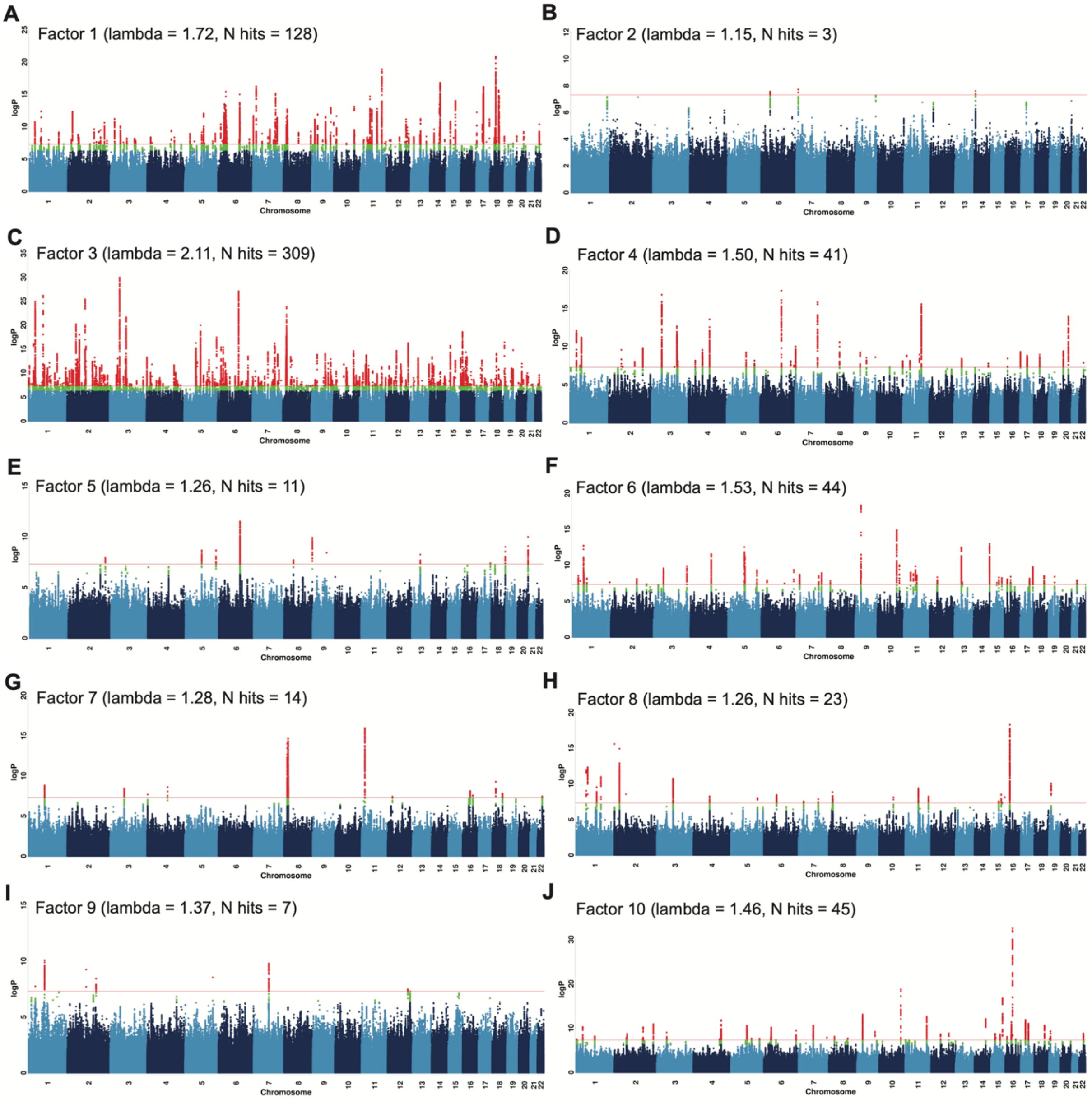


**Supplementary Figure 5**

Manhattan plots of GWAS on top 10 factors from Softimpute; red line shows the genome-wide significance threshold corresponding to P value $5x10^{-8}$.

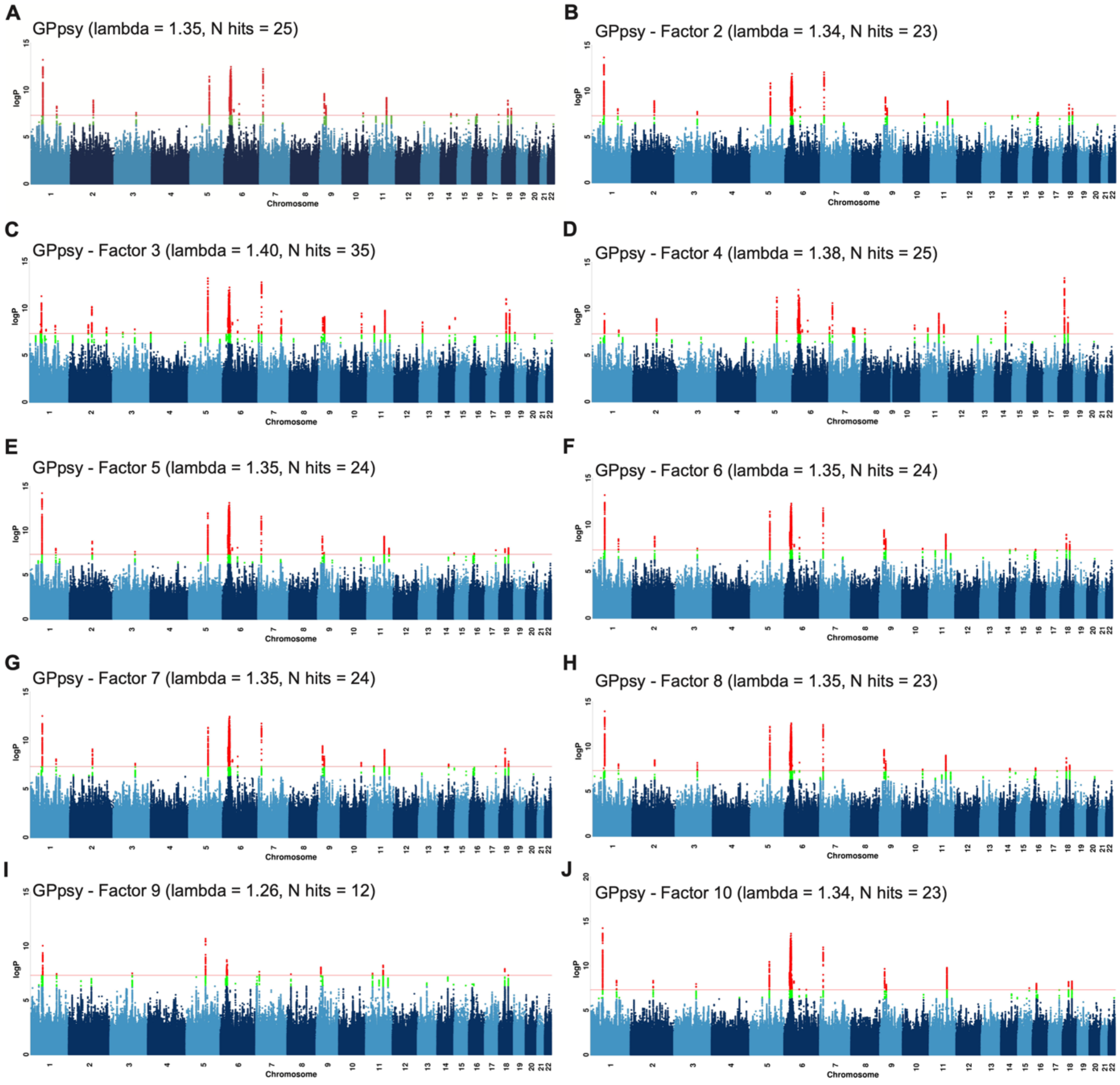


### Supplementary Figure 6

Manhattan plots of GWAS on (A) GPpsy and (B-J) GPpsy regressing out each of the top 9 factors from Softimpute; red line shows the genome-wide significance threshold corresponding to P value $5x10^{-8}$.

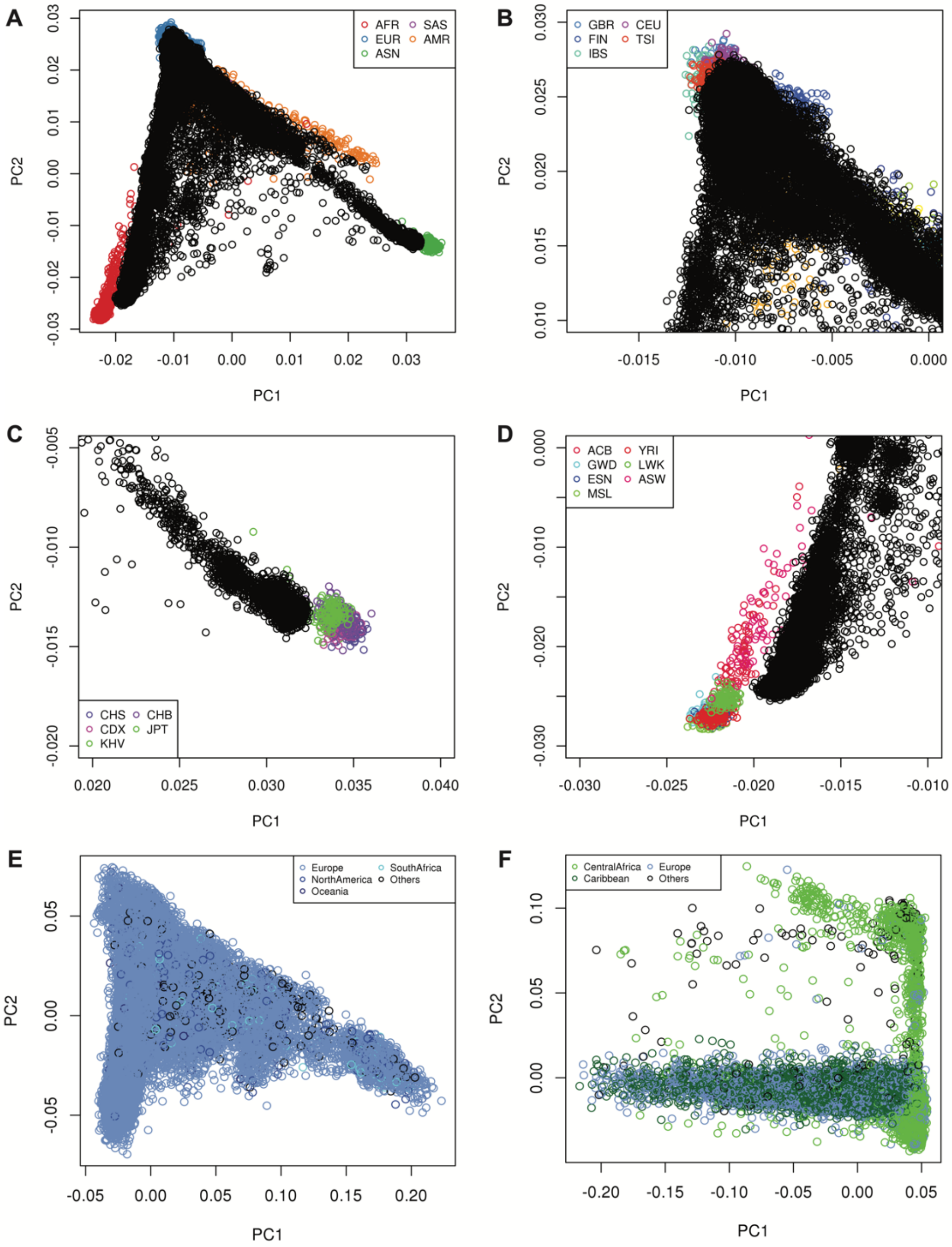


**Supplementary Figure 7**

**(A)** PC1 and PC2 of 150,213 non-White British individuals in UKBiobank projected onto PC1 and PC2 of the 1000G coloured by their super-populations: African (AFR, red), South Asian (SAS, purple), European (EUR, blue), Admixed American (AMR, orange) and Asian (ASN, green); (**B-D)** zoom in of A with 122,710, 8,040 and 2,530 individuals who clustered with the individuals in 1000G with EUR, AFR and ASN ancestries respectively; **(E, F)** PC1 and PC2 of PCA performed only in the EUR and AFR ancestry individuals in UKB, coloured by self-reported ancestries.

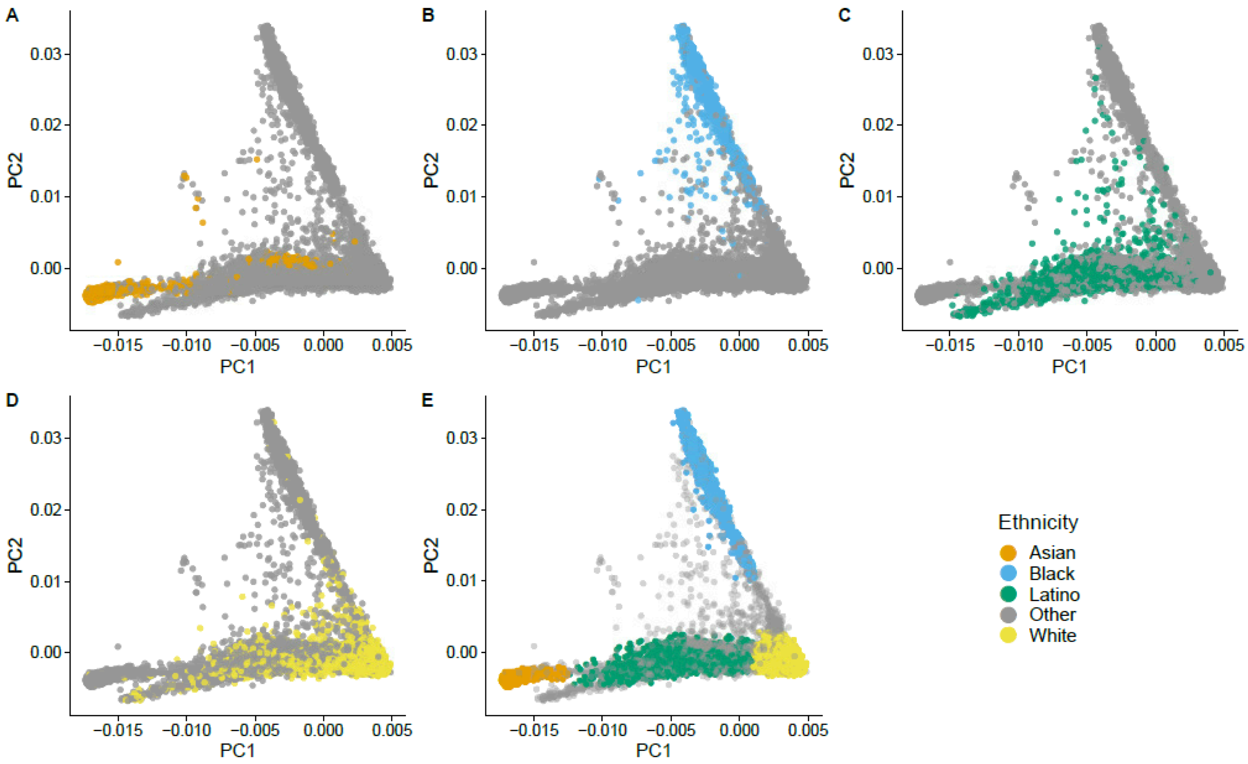


**Supplementary Figure 8**

PC1 and PC2 of individuals in the ATLAS dataset coloured by their reported self-identified race/ethnicity (SIRE), projected onto PC1 and PC2 of the 1000G.

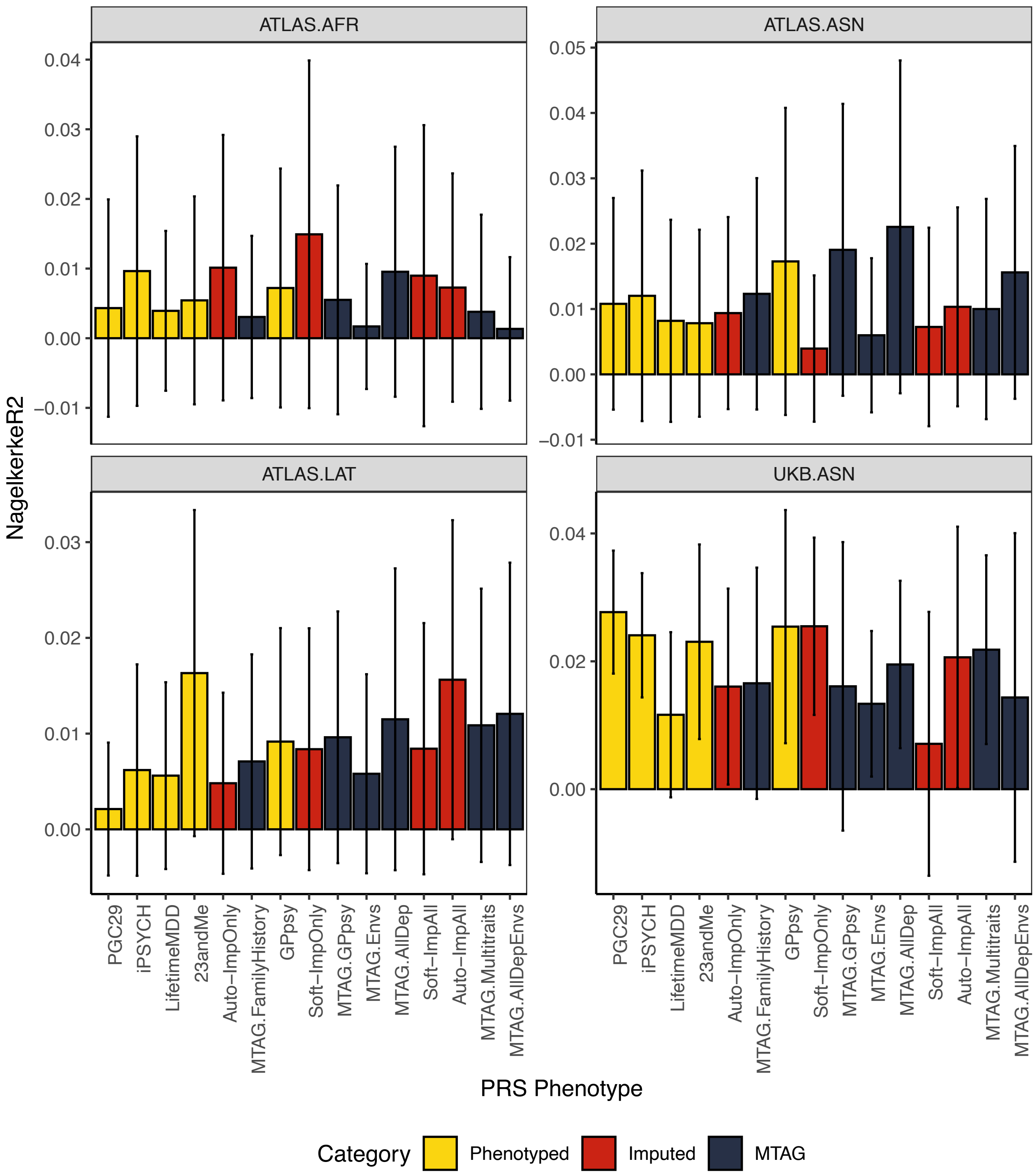


**Supplementary Figure 9**

Out-of-sample PRS prediction accuracy in four additional cohorts of non-European ancestries: individuals who identify as Black (ATLAS.AFR), Asian (ATLAS.ASN), Latino (ATLAS.LAT) in ATLAS, and individuals with East Asian ancestry in UKB.

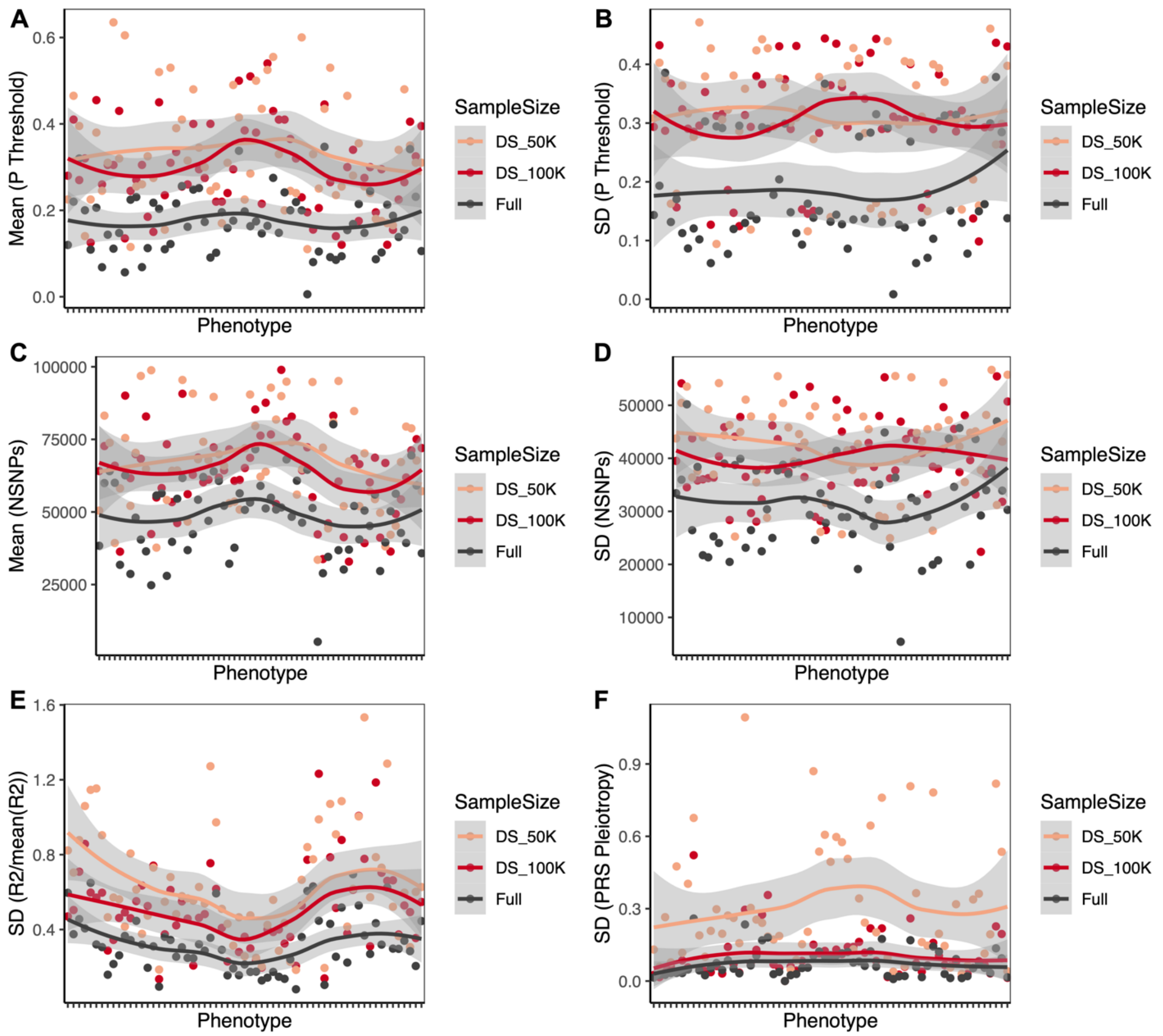


**Supplementary Figure 10**

Effect of training GWAS sample size of GPpsy on **(A)** mean and **(B)** variance in P value threshold, **(C)** mean and **(D)** variance in number of SNPs used in PRS construction, **(E)** coefficient of variation in PRS prediction $R^2$, and **(F)** PRS Pleiotropy. These analyses are performed on 62 phenotypes in UKB (which are significantly predicted by at least one full-sample PRS, as shown in **Figure 6**).

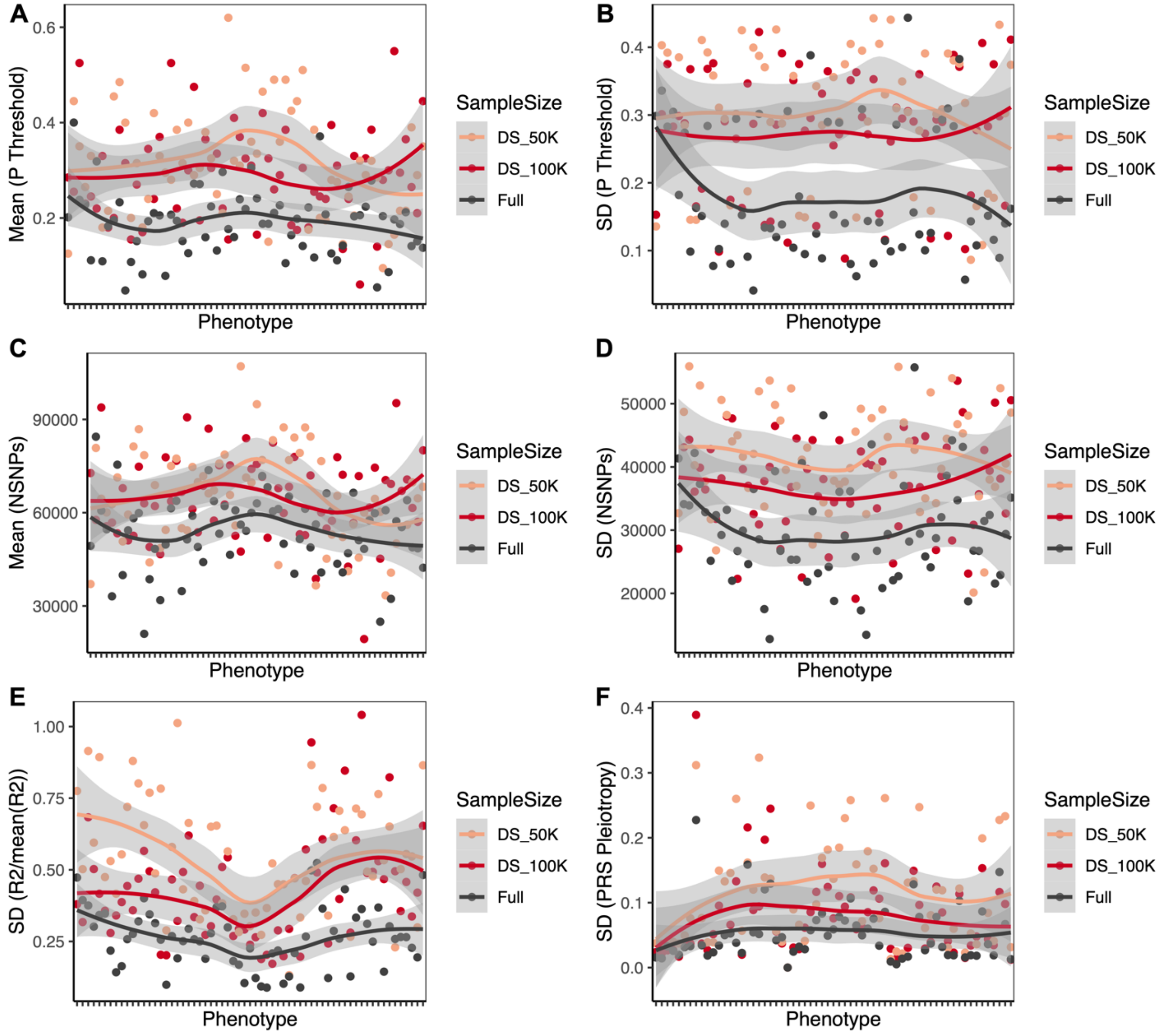


**Supplementary Figure 11**

Effect of training GWAS sample size of Soft-ImpAll on **(A)** mean and **(B)** variance in P value threshold, **(C)** mean and **(D)** variance in number of SNPs used in PRS construction, **(E)** coefficient of variation in PRS prediction $R^2$, and **(F)** PRS Pleiotropy. These analyses are performed on 62 phenotypes in UKB (which are significantly predicted by at least one full-sample PRS, as shown in **Figure 6**).

**Supplementary Tables**

**Supplementary Table 1:** Definitions, data field codes and sample sizes of all MDD-related phenotypes in the UKB that are input for SoftImpute and Autocomplete. Note that phenotypes included in our imputation are not one-to-one with lines in the Table, because some lines correspond to multiple phenotypes (e.g., PC1-20) and some phenotypes correspond to multiple lines (e.g., ICD10 code-based definitions of MDD).

**Supplementary Table 2**: Softimpute R2s of 217 phenotypes in the baseline model (using all 217 phenotypes and all individuals in imputation, including sex, age and 20PCs), in females only, in males only, using phenotypes in MTAG.All (and sex, age and 20PCs) only, and adding in BMI as an extra phenotype.

**Supplementary Table 3:** Genome-wide significant GWAS hits ($P < 5x10^{-8}$, only independent hits are shown) in Softimpute and Autocomplete GWAS on imputed only LifetimeMDD (ImpOnly) and imputed and observed LifetimeMDD combined (ImpAll).

**Supplementary Table 4**: Genome-wide significant GWAS hits ($P < 5x10^{-8}$, only independent hits are shown) from all MTAG runs on LifetimeMDD.

**Supplementary Table 5**: Observed and liability scale SNP-based heritability of all MDD phenotypes used in this study along with the population prevalences (K) used for their calculation; Neff are given as the effective sample size after accounting for case/control imbalance in each observed phenotype; sample sizes N and Neff for MTAG results are identical, given as the effective sample size outputs from MTAG, which refers to the power-equivalent sample sizes for a single phenotype vs multiple phenotypes.

**Supplementary Table 6**: Description, sample sizes, effective sample size (adjusted for case/control numbers) and citations of GWAS on other MDD cohorts referenced and used in this study.

**Supplementary Table 7**: Number of individuals in the UCLA ATLAS study of each reported self-identified race/ethnicity (SIRE), and the number of individuals from each SIRE qualifying for Phecode 296.2 (ATLAS.DPR) and Phecode 296.22 (ATLAS.MDD).

**Supplementary Table 8**: Phecodes in the UCLA ATLAS study used to define MDD cases: 296.2 (ATLAS.DPR) for the superset of depressive illness, corresponding to ICD10 codes for depressive disorders; and Phecode 296.22 (ATLAS.MDD) for the more stringently defined MDD disease status that corresponds to ICD10 codes for Major Depressive Disorder.
